# Inferring macroscopic intrinsic neural timescales using optimal control theory

**DOI:** 10.1101/2025.04.24.650287

**Authors:** Jason Z. Kim, Richard F. Betzel, Ahmad Beyh, Amber Howell, Amy Kuceyeski, Bart Larsen, Caio Seguin, Xi-Han Zhang, Avram Holmes, Linden Parkes

## Abstract

The temporal evolution of brain activity relies on complex interactions within and between brain regions that are mediated by neurobiology and connectivity. To understand these interactions, many large-scale efforts have measured structural connectivity, neural activity, gene expression, and cognition across multiple modalities and species. However, data-driven discovery of large-scale activity models remains difficult owing to the lack of flexible quantitative frameworks for estimating the interplay between brain structure and function while preserving biophysical realism. Here, we provide such a framework by integrating network control theory (NCT) with automatic differentiation to estimate model parameters with greater biophysical realism from data. Specifically, we estimate the structural form of regional self-inhibition—a quantity that is experimentally difficult to measure—from MRI data. Next, we demonstrate that the resulting model-based self-inhibition parameters correlate significantly with regions’ intrinsic neural timescales (INTs), neurobiological measures of gene expression and cell-type densities, as well as behavioral measures of cognition. We demonstrate consistent results across multiple datasets and species. Finally, we demonstrate that our self-inhibition parameters enable the efficient control of brain dynamics from fewer brain regions. Taken together, our results provide a simple and flexible quantitative framework that more accurately captures the interplay between brain structure, function, and dynamics with greater biophysical realism.

## 1 Introduction

The relationship between brain structure and function is underpinned by the complex interplay between microscale cellular processes that govern regional neural dynamics and the macroscale connectivity that binds those dynamics together [1]. For example, variations in the timescales of regions’ intrinsic neural dynamics [2] are governed by diverse gradients of gene expression and cell-type densities that pattern the cortex [3–5]. These gradients also shape the projections between regions [6–8], giving rise to a complex topology of whole-brain structural connectivity known as the *connectome*. In turn, this topology enables large-scale patterns of brain activity [9–11] that support cognition and behavior. Understanding this multi-scale structure-function coupling (SFC) in the brain is a core goal of network neuroscience.

Numerous studies have investigated whole-brain SFC, finding diverse evidence for its multi-scale features [1]. For example, prior work has shown that SFC is stronger between brain regions that (i) are directly connected within the connectome compared to those that are only indirectly connected [12, 13] and (ii) have relatively restricted functional roles (e.g., primary sensory and motor cortex) compared to those that exhibit greater functional diversity (e.g., association cortex) [14, 15]. These effects give rise to a spatial pattern of SFC that varies systematically across the brain, wherein function becomes increasingly untethered from structure in higher-order regions [1, 16]. Such hierarchically-organized SFC has also been observed in the macaque [17] and mouse [18] brains, suggesting that the spatial patterning of SFC is phylogenetically conserved. Complementing these findings, brain regions’ intrinsic neural timescales (INTs)—which are thought to reflect their local computational capacity [2]—are also correlated with this pattern of SFC, as well as with diverse features of macroscale structure and microscale biology [2, 19–22]. For example, in both humans and mice, INTs correlate tightly with the strength of regions’ structural connectivity to the rest of the brain [19, 20], as well as the expression of genes that track diverse types of inhibitory interneurons across the cortex [3, 5]. Together, these findings demonstrate a complex multi-scale relationship between brain structure and function that is conserved across species.

The presence of multi-scale SFC in the brain has motivated the development of a broad range of modeling approaches designed to probe the nature of this relationship. One promising framework is network control theory (NCT) [23–30]. NCT stipulates neural dynamics that evolve on the connectome and models control inputs that guide those dynamics to transition between empirically measured brain states. Here, a *brain state*, ***x***(*t*), describes the level of activity present at each of *N* nodes at various time points, *t*_0_ ≤ *t* ≤ *T*, encoded as an *N* -dimensional vector. Of particular biological relevance is *optimal control*, which finds the exogenous inputs, ***u***(*t*), that control the brain to transition between two activity states while minimizing (i) the magnitude of those inputs (i.e. the *control energy*), and (ii) deviations of the state trajectory from a user-defined reference brain state [23]. Here, *control energy* measures how efficiently brain function can be controlled from brain structure; lower control energy encodes more efficient control and, thus, stronger SFC. Optimal control has been used to study SFC across many neuroscientific contexts [8, 31–39], including neurodevelopment [31], psychiatry [34], neurostimulation [36], and cognition [32]. In each of these contexts, NCT has been able to flexibly probe how brain structure constrains brain function. However, these applications of NCT have relied upon a set of simplifying assumptions that lack biophysical realism, limiting the capacity for NCT to yield multi-scale insights into SFC.

Recently, there has been growing interest in challenging these assumptions with a view to better characterizing multi-scale SFC. For example, prior work has focused on adjusting the strength of the exogenous control inputs to the brain by modifying the input matrix [33, 39–43]. Conventionally, the input matrix comprises a simple one-to-one mapping assumption, wherein *N* control signals are input to *N* nodes with equal weighting [23, 44]. This simple setup asserts that no control input should have more (or less) control of nodes’ dynamics than any other. Recent work has challenged this assumption by modifying the weights of the input matrix to follow spatial patterns of neurobiology [33, 39–42], such as cortical maps of neurotransmitter receptor density. In doing so, these studies have characterized how nodes’ biological properties impact *control energy* across a broad range of state transitions [41]. Though straightforward to implement (but see [23] for caveats), this approach assumes that the best way to parameterize NCT using neurobiology lies in modifying the exogenous control of system dynamics. An alternative approach, that has thus far been unexplored, is to optimize the intrinsic dynamics of the system’s nodes to better represent biology.

Here, we present a simple and flexible extension to the optimal control framework that optimizes the parameters that govern nodes’ intrinsic dynamics. In NCT, each node’s intrinsic dynamics are encoded by an internal decay rate [23], which governs that node’s INTs and can be thought of as its propensity towards self-inhibition; greater self-inhibition yields faster dissipation of perturbations, which equates to faster INTs. Conventionally, decay rates are set uniformly across the connectome (*uniform INTs*). However, this approach is at odds with our understanding of how neuronal time scales vary across the brain in tandem with spatially-patterned properties of neurobiology [2, 19–22]. Thus, to improve the biological realism of NCT, we developed a data-driven approach to optimizing nodes’ internal decay rates. Specifically, we implement learnable decay rates that minimize the distance between the model’s controlled state trajectory and an empirical reference state, thereby yielding *optimized INTs*. Compared to previous work that applied variable weights to the input matrix [33, 39–42], our approach does not require *a priori* knowledge of neurobiology. Rather, our approach assumes that the effects of neurobiology are already encoded in the connectome and empirically measured states, and thus optimizing decay rates will uncover INTs that reflect underlying neurobiology. In turn, this approach allows us to both validate optimized INTs against known neurobiological correlates, and to fit our model on a per-transition and a per-subject basis. Furthermore, our approach can be combined with preexisting methods for modifying the input matrix, including our own recently developed approaches [43].

Using our approach, we find that using optimized decay rates yields lower control energy associated with transitions between empirical brain states, demonstrating stronger SFC. We also find that optimized decay rates correlate with *in vivo* empirical measures of INTs as well as *ex vivo* measures of gene expression [45] and cell-type densities [46] that underpin SFC in humans. Additionally, we find that these correlations with neurobiology were conserved in the mouse connectome [47, 48], demonstrating convergence across species. Finally, when applied to single-subject human connectomes, we find that our approach significantly improves the out-of-sample prediction of behavior in healthy young adults. In sum, we provide a novel extension to NCT to better model and validate the complex multi-scale interaction between brain structure and function with subject-level specificity.

## 2 Results

Here, we present an overview of our approach to modeling and comparing *uniform* and *optimized* neural dynamics using network control theory (NCT). We develop and apply our approach to multi-modal neuroimaging data taken from the Human Connectome Project Young Adult (HCP-YA) sample [49]. We replicate our findings in two separate datasets: (i) another human dataset called Microstructure-Informed Connectomics (MICA-MICs) [50] as well as (ii) data from the mouse brain taken from the Allen Mouse Brain Connectivity Atlas (AHBA) [47, 48]. From the human datasets, we extracted group-averaged structural connectomes using dMRI and regional neural dynamics using resting-state fMRI (rs-fMRI) (see Materials and Methods).

### 2.1 Network control theory: optimizing internal decay rates

The use of NCT in neuroscience is well-established and we refer readers to our recent protocol paper [23] for extended discussion on its definition and application to connectome data [51, 52]. Here, we used *optimal control theory* [30, 44], defined using a continuous-time system, to simulate neural dynamics unfolding upon the structural connectome and to study transitions between brain states extracted from rs-fMRI. Optimal control employs a time-varying input, ***u***(*t*), to drive neural activity, ***x***(*t*), from an initial state, ***x***_0_, to a target state, ***x***_*T*_, while balancing constraints on both the magnitude of ***u***(*t*) and ***x***(*t*) (Fig. 1A, B). The magnitude of ***u***(*t*) represents the control energy, *E*, a measure of how much simulated “effort” is required to complete a given state transition. Critical to the current study, however, is the magnitude of ***x***(*t*). In optimal control, the magnitude of ***x***(*t*) is penalized for departing too much from a user-defined reference state, ***x***_*r*_. Here, we set ***x***_*r*_ to ***x***_*T*_, which results in neural activity being constrained toward the target state throughout a transition. While prior work has examined the impact of removing the state magnitude constraint (i.e., *minimum control energy*) [28], so far no study has sought to optimize this constraint by modifying the connectivity to minimize the delta between ***x***(*t*) and ***x***_*r*_. Here, we extend NCT to allow for this minimization.

**Figure 1.**
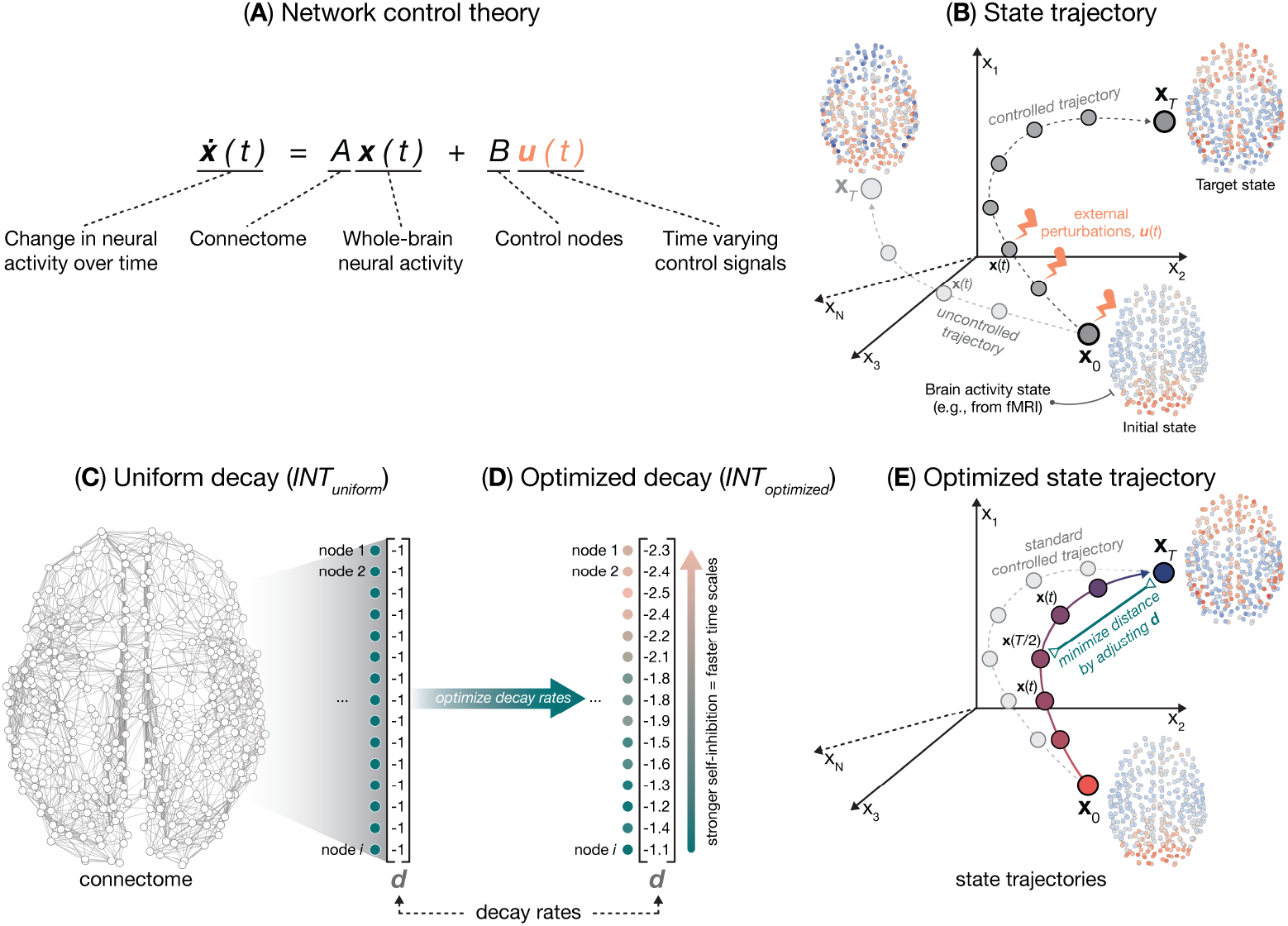
Illustration of network control framework using optimized decay rates. (**A**) In network control theory (NCT), the human connectome is used to model controlled neural dynamics according to 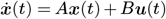.(**B**) Using this equation, NCT allows researchers to model the control signals, ***u***(*t*), that guide neural activity, ***x***(*t*), to transition between an initial state, ***x***_0_, and a target state, ***x***_*T*_ . (**C**) Within NCT, nodes’ local dynamics are parameterized by their internal decay rates, ***d***. These decay rates determine nodes’ tendency to inhibit their own dynamics; stronger negative values correspond to stronger self-inhibition and thus faster intrinsic neural timescales (INTs). Typically, decay rates are set uniformly across the brain to a value of ***d*** = −1, as shown here. (**D**) Here, we develop an approach to optimizing nodes’ internal decay rates that allows them to vary across the brain, giving rise to heterogeneous INTs. (**E**) Optimized decay rates are found by minimizing the distance between ***x***(*t*) and a reference state, ***x***_*r*_, at the midpoint of the state transition, ***x***(*T/*2). Here, ***x***_*r*_ = ***x***_*T*_ . As a result, we uncover ***d*** that minimizes the distance between ***x***(*t*) and ***x***_*T*_ over time. We achieve this minimization by treating ***d*** as a free parameter in a gradient descent algorithm implemented in *PyTorch* (see Materials and Methods).

Our extension to NCT involves modifying the decay rates, ***d***, of nodes’ internal dynamics (see Materials and Methods) (Fig. 1C-E). In NCT, prior to simulating a state transition, each node’s intrinsic dynamics is set to decay over time in the absence of external input, which ensures system stability [23, 28]. In a continuous-time system, this internal decay is set uniformly across nodes by subtracting the identity matrix (*I*) from the connectome (*A*) during normalization (see Materials and Methods). This approach establishes a baseline decay rate of *d* = −1 for each node, which we refer to as *uniform INTs* (Fig. 1C; *INT*_uniform_). Here, we optimize these decay rates to minimize the distance between the controlled state trajectory and the reference state. This approach is data-driven and results in a set of heterogenous decay rates that give rise to variable INTs across the brain; an approach we refer to as *optimized INTs* (Fig. 1D; *INT*_optimized_). For a given state transition, we achieved this optimization by performing gradient descent *via* automatic differentiation on the difference between the controlled state trajectory, ***x***(*t*), and the reference state, ***x***_*r*_, while adjusting ***d*** (Fig. 1E). Once converged, this algorithm provides a unique decay rate for each node and for each state transition. Then, these optimized decay rates can be (i) used to calculate control energy for their associated transition and (ii) validated against patterns of neurobiology known to influence empirical connectivity and neural dynamics.

### 2.2 Optimizing regions’ internal dynamics leads to reduced control energy

Using the above approach, we trained our algorithm to complete transitions between seven pairs of brain states that each represent resting-state coactivation patterns (CAPs; Fig. 2A). Following prior work [32, 39, 42], we extracted these brain states by applying *k*-means clustering to rs-fMRI time series data (see Materials and Methods). We chose seven clusters because the difference in the inertia between *k* and *k* + 1 suggested that this solution was optimal (Fig. S1A; see Fig. S1B for the correlations between brain states and see Supplementary Results for sensitivity analysis spanning *k* = 2 − 13). As expected, we found that our seven brain states aligned broadly with previously reported canonical CAPs [32, 39] (see Fig S1D for overlap between brain states and canonical brain systems [53, 54]). Across all pairs of state transitions, we plot the mean and standard deviation of the optimized decay rates in brain space, and find that they are spatially-patterned (Fig. 2B, see Fig. S3 for transition-specific optimized decay rates).

**Figure 2.**
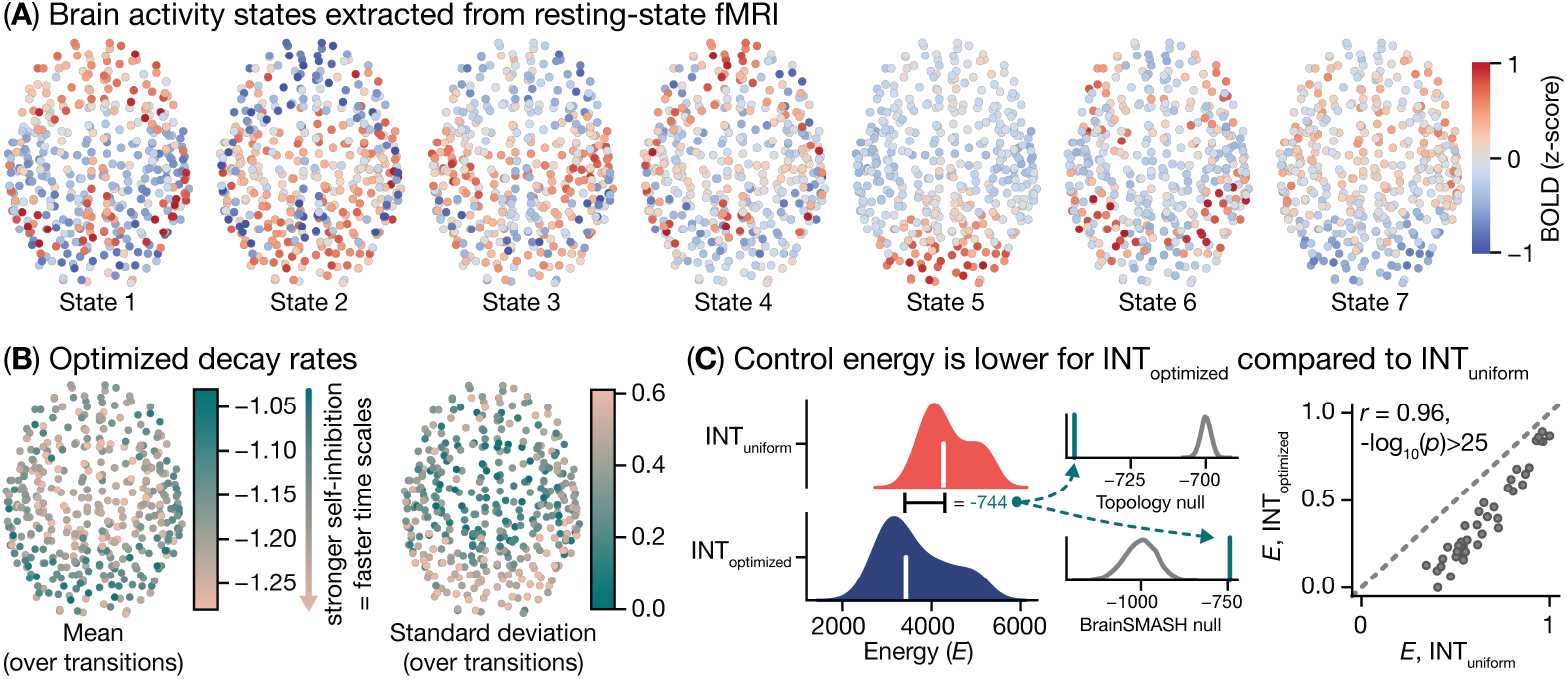
Optimizing nodes’ intrinsic neural time scales results in lower control energy for brain state transitions. (**A**) Spatial topography of seven empirically-derived brain states. These states were extracted using *k*-means clustering applied to resting-state fMRI time series data; see Fig S1 for further characterization of these brain states. We used these brain states to perform NCT analyses and optimize our model. (**B**) Fully-trained regional optimized internal decay rates. Here, the mean (left) and standard deviations (right) were estimated over 42 state transitions. For the mean, more negative values correspond to stronger self-inhibition and thus faster intrinsic neural time scales (INTs; see arrow). (**C**) Control energy associated with each state transition plotted for our *INT*_optimized_ model compared to the standard *INT*_uniform_ model.

Our *INT*_optimized_ model is designed to increase the biophysical realism of NCT by modeling biological variation in the decay rates. This increased realism should more accurately reflect natural brain dynamics, which we predict will lead to stronger SFC and lower control energy. To examine this prediction, once our algorithm was fully trained (see Fig. S2), we calculated control energy, *E*, separately using both our *INT*_uniform_ and *INT*_optimized_ models. To control for differences in the overall level of self-inhibition between the two models, we set all values of ***d*** for the *INT*_uniform_ model to the mean of ***d*** from the *INT*_optimized_ model 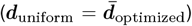.As predicted, we found that control energy was lower for *INT*_optimized_ compared to *INT*_uniform_ (Fig. 2C, left; 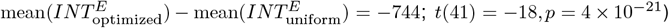. Furthermore, this reduction in control energy was highly consistent across state transitions (Fig. 2C, right; *r* = 0.96, *p* = 2 *×* 10^−24^). Critically, these effects are not-trivial because the decay rates are optimized to minimize the distance between the controlled state trajectory and a reference state (∥***x***(*T/*2) − ***x***_*T*_ ∥), and *not* the control energy. These results demonstrate that our extension to NCT allows for more efficient control of large-scale neural dynamics from connectome topology, which in turn narrows the gap between brain structure and function.

To examine whether optimized decay rates conferred a significant energetic advantage specifically for real biological connectivity and activity, we compared the reduction in mean energy (see Fig. 2G) against the following pair of null models [55]. First, to quantify the influence of connectome topology on the energy reduction, we randomly rewired the connectome 5,000 times subject to constraints on nodes’ spatial embedding and the network’s strength distribution [56], and for each we retrained our *INT*_optimized_ model and recomputed the difference of means in control energy. Second, to quantify the influence of the specific state transition on energy reduction, we randomly permuted the values of the target states—and thus the reference states—while preserving the spatial autocorrelation of the fMRI data (brainSMASH [57]), and for each we retrained our *INT*_optimized_ model and recomputed the difference of means in control energy. We found that the magnitude of the energy reduction was significantly larger than expected under the topology null (Fig. 2C, Topology null) and significantly smaller than expected under the brainSMASH null (Fig. 2C, BrainSMASH null). These results provide robust evidence that our optimized decay rates are jointly influenced by the topology of the connectome and the spatial patterning of the target states; as opposed to being driven exclusively by one or the other. Critically, the opposing null effects reported here show that the impact of the connectome and the brain states on the optimized decay rates are in tension with one another, and that our model balances these competing influences throughout optimization. In turn, this finding suggests that while our model gives rise to regional dynamics that are rooted in intrinsic brain activity (i.e., those generated by the connectome), dynamics are able to flexibly update to meet the needs of a specific state transition. Thus, our model-based INTs have both a stable and a flexible component. Together, these results demonstrate our model-based INTs emerge as a product of jointly optimizing on real biological connectivity and activity.

### 2.3 Optimizing regions’ internal dynamics reduces the dimensionality of control signals

Above, we showed that optimizing regions’ intrinsic neural dynamics systematically reduced the control energy associated with transitioning between functional brain states, indicating tighter coupling between structure and function. We observed these results using a *uniform full control set* [23], wherein all nodes of the system, *N*_*system*_, were designated as control nodes, yielding as many control signals as system nodes. However, as documented in our recent work [43], we know that a full control set produces highly correlated—and thus redundant—control signals. We also know that such a control strategy is inconsistent with real-world approaches to modulating brain activity; for example, techniques such as transcranial magnetic stimulation input control signals focally to the brain rather than to all regions at once. As such, we recently examined whether this control signal redundancy could be leveraged to complete state transitions efficiently using only a subset of unique control signals, *N*_*unique*_, delivered to *N*_*system*_ control nodes, where *N*_*unique*_ *<< N*_*system*_ [43]. We found that we could obtain low-energy approximations of optimal control by driving multiple regions with the same control input, demonstrating that control signals are highly compressible. However, this approach still delivered control signals to every system node, leaving open the following question: can the connectome can be controlled using only a subset of control signals delivered to a subset of system nodes? Addressing this question would bring NCT closer to real-world applications of exogenous control. Thus, here we examined whether our *INT*_optimized_ model could complete state transitions using only a subset of system nodes [23].

For each state transition, we computed control energy for both the *INT*_uniform_ and *INT*_optimized_ models using a partial control set [23] (see Materials and Methods). A partial control set is defined by designating only a subset, *k*, of system nodes as control nodes; the smallest possible partial control set comprises only a single control node (*k* = 1), whereas the largest partial control set comprises all but one system nodes (*k* = *N*_*system*_ − 1). We computed control energy over a range of *k* values, each time designating *k* random nodes as control nodes. For each *k*, we repeated this process 20 times, with different randomly sampled control nodes. We found that state transitions completed successfully for both the *INT*_uniform_ and *INT*_optimized_ models with relatively few control nodes (Fig. 3). Specifically, for both models, we observed correlations between ***x***_*T*_ and ***x***(*T*) that were *>* 0.9 (Fig. 3A), and reconstruction errors *<* 1 *×* 10^−2^ (Fig. 3B), when *k* ≥ 124. This results indicates that state transitions completed successfully using control sets that were only ≈ 1*/*3 the size of our system (here, *N*_*system*_ = 400). Critically, the *INT*_optimized_ model yielded this same performance when *k* ≥ 64, indicating that our model achieved successful transitions with control sets that were ≈ 48% smaller than those required for the *INT*_uniform_ model. Furthermore, smaller control sets also produced lower control energy for the *INT*_optimized_ model compared to the *INT*_uniform_ model (Fig. 3C). Thus, the optimized decay rates in our model allowed us to control connectome dynamics both more efficiently (i.e., with fewer nodes) and more easily (i.e., lower control energy) from a partial control set. These results indicate that our approach—which seeks to improve the biophysical realism of NCT—enables the successful control of brain dynamics from relatively focal inputs. In turn, this improvement to model fitting reduces the gap between NCT and real-world applications of brain stimulation.

**Figure 3.**
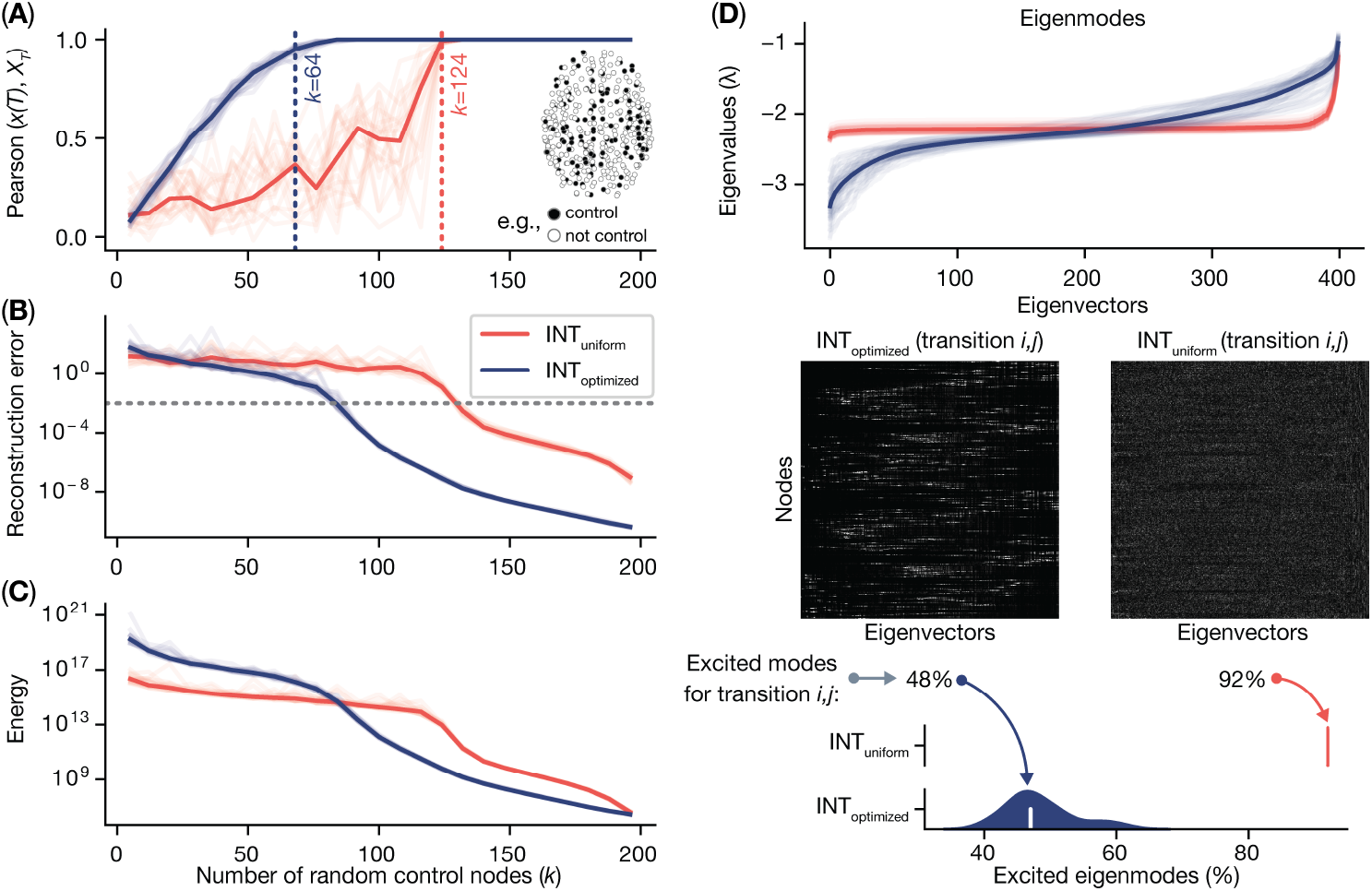
Optimizing nodes’ intrinsic neural time scales results in more efficient control of connectome dynamics from fewer control nodes. We estimated control energy using random partial control sets of increasing size, *k*. For each *k*, we generated 20 random control sets and calculated control energy for all state transitions. (**A**) The correlation between ***x***(*t*) and ***x***_*T*_ for both the *INT*_optimized_ and *INT*_uniform_ models averaged over the 20 control sets at each *k*. (**B**) The reconstruction error for both models averaged over the 20 control sets at each *k*. (**C**) The control energy for both models averaged over the 20 control sets at each *k*. Here, control energy was rescaled by multiplying by *dt*, which was set to 0.001 in our simulations. Together, these plots show that state transitions were completed more efficiently (i.e., with fewer control nodes) and more easily (i.e., lower control energy) when using the *INT*_optimized_ model. (**D**) The connectome eigenmodes for both the *INT*_optimized_ and *INT*_uniform_ models. In the top plot, the eigenvalues for each eigenvector are shown averaged over state transitions. This data shows that the eigenvectors of the system have more diverse eigenvalues when using the *INT*_optimized_ model. In the bottom plots, the corresponding eigenvector matrices are shown for an example transition (middle), alongside their density averaged over state transitions (bottom). This data shows that the nodes of the connectome innervate different subsets of eigenmodes in the *INT*_optimized_ model compared to the *INT*_uniform_ model. On average, nodes innervated only 48% of the eigenmodes for the *INT*_optimized_ model. By contrast, nodes innervated 92% of the eigenmodes for the *INT*_uniform_ model. Thus, under the *INT*_optimized_ model, each node innervates only a relatively sparse set of eigenmodes that exert a more diverse impact on whole-brain dynamics over time, and this leads to more efficient control over state transitions.

To provide intuition for the above effects, we performed eigenmode decomposition on the connectome, wherein the diagonals were set either to the transition-specific decay rates from *INT*_optimized_ model or to their mean. This analysis revealed that the above effects (see Fig. 3A-C) were driven by the *INT*_optimized_ model giving rise to more diverse whole-brain dynamics (Fig. 3D). Specifically, compared to the *INT*_uniform_ model, the eigenmodes of the *INT*_optimized_ model produced a wider ranges of eigenvalues across all state transitions (Fig. 3D, top), indicating greater variability in how they influence whole-brain dynamics over time. This variability strongly impacts controllability when *N*_*unique*_ *< N*_*system*_ because there are not enough unique control inputs to independently drive a successful transition for each node [58]. Hence, successful transitions must be achieved by relying on the differential response of the eigenmodes across time to pattern a temporal sequence of *N*_*unique*_ inputs which, through the differential response of eigenmodes, enables successful control with fewer nodes.

Additionally, the corresponding eigenvectors were noticeably more sparse for the *INT*_optimized_ model compared to those from the *INT*_uniform_ model (Fig. 3D, bottom). This result indicates that single nodes innervate only relatively small subsets of the eigenmodes in the *INT*_optimized_ model; nodes innervated, on average, ∼ 48% of the eigenmodes for the *INT*_optimized_ model compared to 92% for the *INT*_uniform_ model. Hence, our *INT*_optimized_ model gives rise to a greater ability to control targeted subsets of dynamical modes from single nodes. Together, these results show that, under the *INT*_uniform_ model, all nodes of the connectome tend to innervate the majority of the eigenmodes of the system, and that the effect of these modes on whole-brain dynamics are largely consistent over time (i.e., homogenous eigenvalues). By contrast, under the *INT*_optimized_ model, each node innervates only a sparse set of eigenmodes that exert a more diverse impact on whole-brain dynamics over time (i.e., heterogenous eigenvalues). In turn, this difference leads to the efficient completion of state transitions from fewer control nodes.

### 2.4 Optimized dynamics correlate with regions’ intrinsic neural time scales

Having demonstrated that introducing variable self-inhibition to connectome dynamics yielded a consistent reduction in control energy, we next sought to characterize the variability of our optimized decay rates across state transitions as well as validate them against empirical measures of intrinsic neuronal time scales [2]. Unlike weighting the input matrix (*B*) *a priori* as in previous work [33, 39–43], our approach involves optimizing nodes’ intrinsic dynamics on a per-transition basis. In turn, this process gives rise to a vector of optimized decay rates for each transition that encodes a spatial pattern of heterogenous self-inhibition across the brain. To develop intuition for our optimized decay rates, we initially plotted them averaged over state transitions above in Fig. 2F. Here, using principal component analysis (PCA), we extracted the two dominant sources of variance in optimized decay rates across state transitions (see Materials and Methods). PC 1 and PC 2, which accounted for 35% and 22% of the variance, respectively, are shown in Fig. 4 (see Fig. S4 for plots of PCs 3-5 as well as correspondence between PCs 1-5 and canonical brain systems). For each of these PCs, greater scores correspond to slower INTs in our model across diverse transitions.

**Figure 4.**
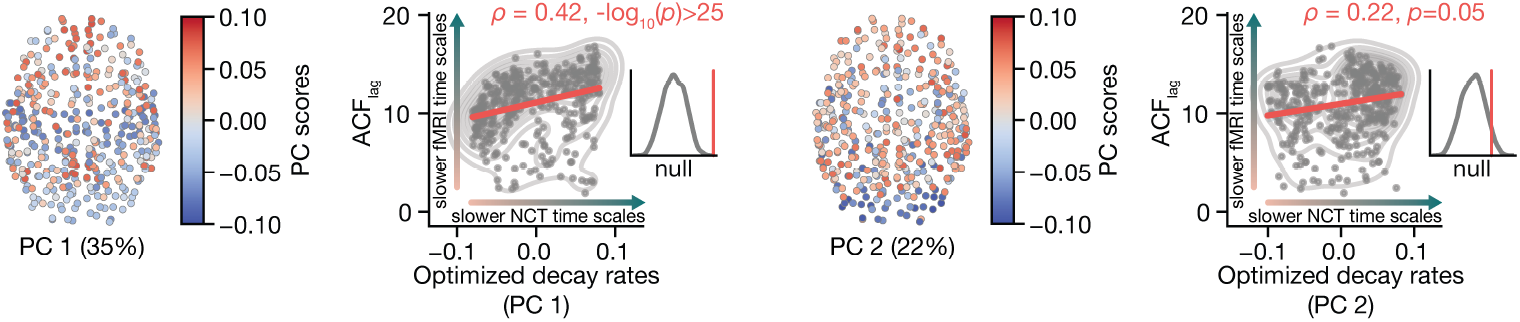
Model-based intrinsic neural time scales (INTs) correlate with empirically-measured INTs. We used principal component analysis (PCA) to extract topographic maps that described the two dominant sources of variance in optimized decay rates across state transitions. Here, higher scores on PC 1 and PC 2 corresponded to slower time scales in our model based on correlating each PC with the mean of the optimized decay rates (taken from Fig. 2B). Next, we measured empirical INTs by fitting an autocorrelation function (ACF) to nodes’ rs-fMRI time series data and counting the number of frames it took for the ACF to decay to 0 (*ACF*_lag_). Higher *ACF*_lag_ values corresponded to slower decay and therefore slower empirical INTs. Both PC 1 and PC 2 correlated with positively with *ACF*_lag_, indicating consistency between model-based and empirically-measured INTs. We assigned p-values to these correlations using brainSMASH

We correlated this compact representation of optimized decay rates with an empirical measure of INTs. Specifically, we calculated the decay lag of the autocorrelation coefficient of rs-fMRI time series (*ACF*_lag_; see Materials and Methods). Here, higher values indicate slower decay of the ACF and thus slower empirically-measured INTs. Then, we compared the correlation between each PC and *ACF*_lag_ against a null model derived from brainSMASH [57]. Here, brainSMASH was used to generate spatial autocorrelation-preserving surrogates of *ACF*_lag_ (see Materials and Methods). We found that *ACF*_lag_ correlated positively with PC 1 (Fig. 4, left; *ρ* = 0.44, −*log*_10_(*p*_*smash*_) *>* 25) as well as PC 2 (Fig. 4, right; *ρ* = 0.22, *p*_*smash*_ = 0.05), albeit to a lesser extent. Note, we found that the mean of optimized decay rates (taken from Fig. 2F) also correlated positively with *ACF*_lag_ (*ρ* = 0.70, −*log*_10_(*p*_*smash*_) *>* 25). These results illustrate that the two dominant sources of variance in our optimized decay rates each coupled to empirical INTs. Critically, this finding shows that optimized decay rates from NCT are representative of INTs derived from real data. As discussed previously (see Fig. 2F), higher (i.e., less negative) decay rates correspond to weaker self-inhibition, which gives rise to slower model-based dynamics. Thus, brain regions with slower dynamics in our model (higher scores on PC 1 and PC 2) also exhibit slower dynamics measured using rs-fMRI. Note, we also correlated PC 1, PC 2, as well as the mean of optimized decay rates with regional degree and strength estimated from the structural connectome and found no significant relationships (all *ps >* 0.05). The largest effect was a small positive correlation between PC 1 and strength (*ρ* = 0.11, *p* = 0.06). Thus, our optimized decay rates do not simply recapitulate lower-order topological properties of the connectome.

### 2.5 Optimized dynamics correlate with measures of neurobiology that underpin neural dynamics

Above, we showed that the dynamics from our *INT*_optimized_ model correlated with empirical neural dynamics measured using rs-fMRI. This analysis validated our model’s simulated dynamics empirically at the macroscale, but what about the microscale features of neurobiology that are thought to underpin brain dynamics? Here, we investigate this question by comparing optimized decay rates with maps of whole-brain gene expression [45, 59–61] and cell-type densities [46, 62] (see Materials and Methods), as well as intracortical myelin content [63].

Recent work has demonstrated that whole-brain connectivity and dynamics may be underpinned by diverse features of gene expression measured at the cellular level [4, 64–67]. For example, genes coding for the concentration of inhibitory interneurons across the cortex correlate tightly with cortical gradients of INTs [2, 3, 22] and functional connectivity [46]; together, these features follow a cortical axis of variation that spans primary sensorimotor regions to higher-order association regions [3, 4, 68, 69]. As such, leveraging the Allen Institute Human Brain Atlas (AHBA) [45], we extracted parcellated maps of gene transcripts [59] and correlated those maps with our optimized decay rates (see Materials and Methods). Additionally, drawing on our recent work [46], we analyzed parcellated maps of imputed cell-type densities; these latter maps provide more subtle insight into spatial patterns of neurobiology by teasing apart different cell-types that express similar genes. Finally, prior work has shown that intracortical myelin content measured using the ratio of T1w/T2w imaging represents an *in vivo* proxy for AHBA gene expression [4]. Thus, we also compared our optimized decay rates against intracortical myelin.

First, we examined the coupling between PC 1 and PC 2 of optimized decay rates and maps of somatostatin (SST) and parvalbumin (PVALB) genes. SST and PVALB are anti-correlated across the cortex [3] and track distinct sub-types of inhibitory interneurons that differ in terms of their neuronal cell targeting [70–72], their developmental timing [73, 74], and their control over neural dynamics [73, 75]. For example, SST-expressing cells synapse onto the dendrites of pyramidal neurons, which positions them to gate incoming signals before they reach the soma. By contrast, PVALB-expressing cells regulate the output firing of pyramidal neurons and fall into one of two categories: (i) PVALB-Basket cells that synapse perisomatically with pyramidal neurons, and (ii) PVALB-Chandelier cells that synapse with the axon initial segment. These distinct synaptic patterns are thought to endow SST and PVALB interneurons with different inhibitory roles, which in turn give rise to different patterns of neural dynamics across the brain. Consistent with this idea, previous research has shown that higher-order association regions exhibit relatively high SST expression and slower INTs while primary sensorimotor regions exhibit relatively high PVALB expression and faster INTs [3]. In accordance with this literature, using AHBA gene maps, we found that PC 1 of optimized decay rates correlated positively with SST (Fig. 5A, left; *r* = 0.30, *p*_*smash*_ = 3 *×* 10^−2^) and negatively with PVALB (Fig. 5A, right; *ρ* = −0.34, *p*_*smash*_ = 2 *×* 10^−3^). Thus, relatively slow INTs in our model linked to high expression of genes associated with slower neural time scales (SST), while relatively fast INTs in our model linked to high expression of genes associated with faster neural time scales (PVALB). Furthermore, using our imputed cell-type maps [46], we also found that optimized decay rates correlated negatively with PVALB-Basket cells (Fig. 5B, left; *ρ* = −0.25, *p*_*smash*_ = 7 *×* 10^−3^) and positively with PVALB-Chandelier cells (Fig. 5B, right; *ρ* = 0.19, *p*_*smash*_ = 3 *×* 10^−2^). These results suggest that the negative correlation observed for AHBA-PVALB genes may reflect coupling with basket cells rather than chandelier cells. Critically, we could not uncover these relationships using the AHBA alone. Next, we observed a negative correlation between PC 1 and intracortical myelin content (Fig S5; *ρ* = −0.54, −*log*_10_(*p*_*smash*_) *>* 25), which is consistent with the observation that highly myelinated sensorimotor regions exhibit faster INTs [2]. Finally, we observed no significant correlations between PC 2 of optimized decay rates and any of the above brain maps. Thus, while there are two dominant sources of variance in our model-based INTs that correlate with empirical INTs (see Fig. 4), only the first source of variance connects with maps of neurobiology that reflect cortical inhibition. Collectively, these results demonstrate that parameters from our *INT*_optimized_ model track the divergent spatial patterning of distinct inhibitory interneurons across the cortex.

**Figure 5.**
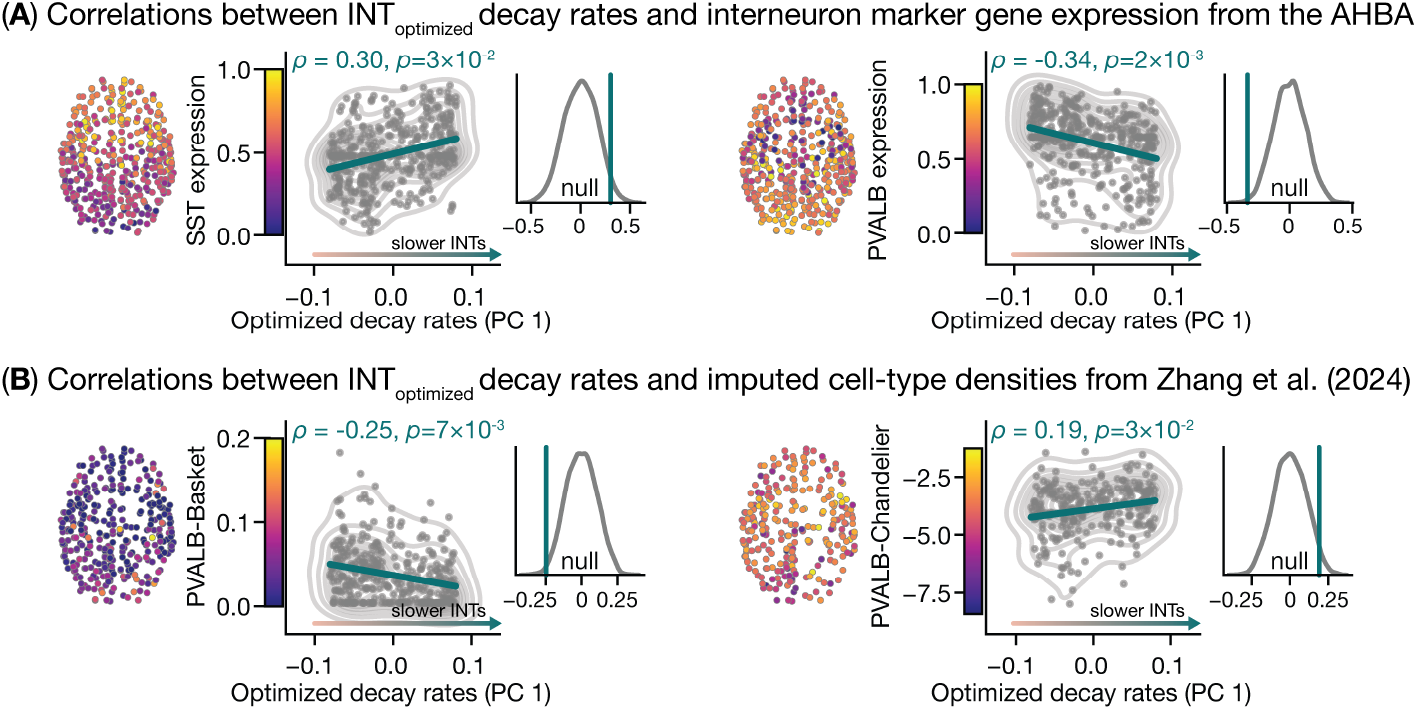
Model-based intrinsic neural time scales correlate with whole-brain maps of inhibitory interneurons. (**A**) Correlations between PC 1 of optimized decay rates (taken from Fig. 4) and whole-brain maps of gene expression extracted from the Allen Human Brain Atlas [45]. We correlated PC 1 with maps of somatostatin (SST, left) and parvalbumin (PVALB, right) genes. PC 1 correlated positively with SST genes and negatively with PVALB genes. We assigned p-values to these correlations using brainSMASH [57]. (**B**) Correlations between PC 1 of optimized decay rates and whole-brain maps of cell-type densities taken from [46]. We correlated PC 1 with cell-type density maps of PVALB-expressing basket cells (left) and PVALB-expressing chandelier cells (right). PC 1 correlated negatively with the former and positively with the latter. We assigned p-values to these correlations using brainSMASH [57].

### 2.6 Results replicate across datasets and species

In this section, we examined whether our model replicated in a second human dataset and generalized across species. Specifically, we replicated our key results using an undirected human connectome taken from the MICA-MICs dataset [50] as well as a directed mouse connectome taken from the Allen Mouse Brain Connectivity Atlas [47, 48] (see Materials and Methods).

When fitting the *INT*_*optimized*_ model to the MICA-MICs dataset, we observed results that were highly consistent with those observed for the HCP-YA dataset. First, we found lower control energy for the *INT*_*optimized*_ model that exhibited the same *tug of war* effect under our twin null models (Fig. 6A, left). Second, we found that the dominant source of variance in optimized decay rates correlated positively with *ACF*_lag_ estimated from fMRI data (Fig. 6A, top right; *ρ* = 0.22, *p*_*smash*_ = 8 *×* 10^−3^) and negatively with *in vivo* microstructure taken from [50] (Fig. 6A, top right; *ρ* = 0.22, *p*_*smash*_ = 8 *×* 10^−3^) (see Materials and Methods). Finally, we observed the same pattern of correlations between optimized decay rates and our gene and cell-type maps. That is, optimized decay rates correlated positively with AHBA-SST, AHBA-PVALB, and PVALB-Chandelier, and negatively with PVALB-Basket (Fig. 6A, bottom).

**Figure 6.**
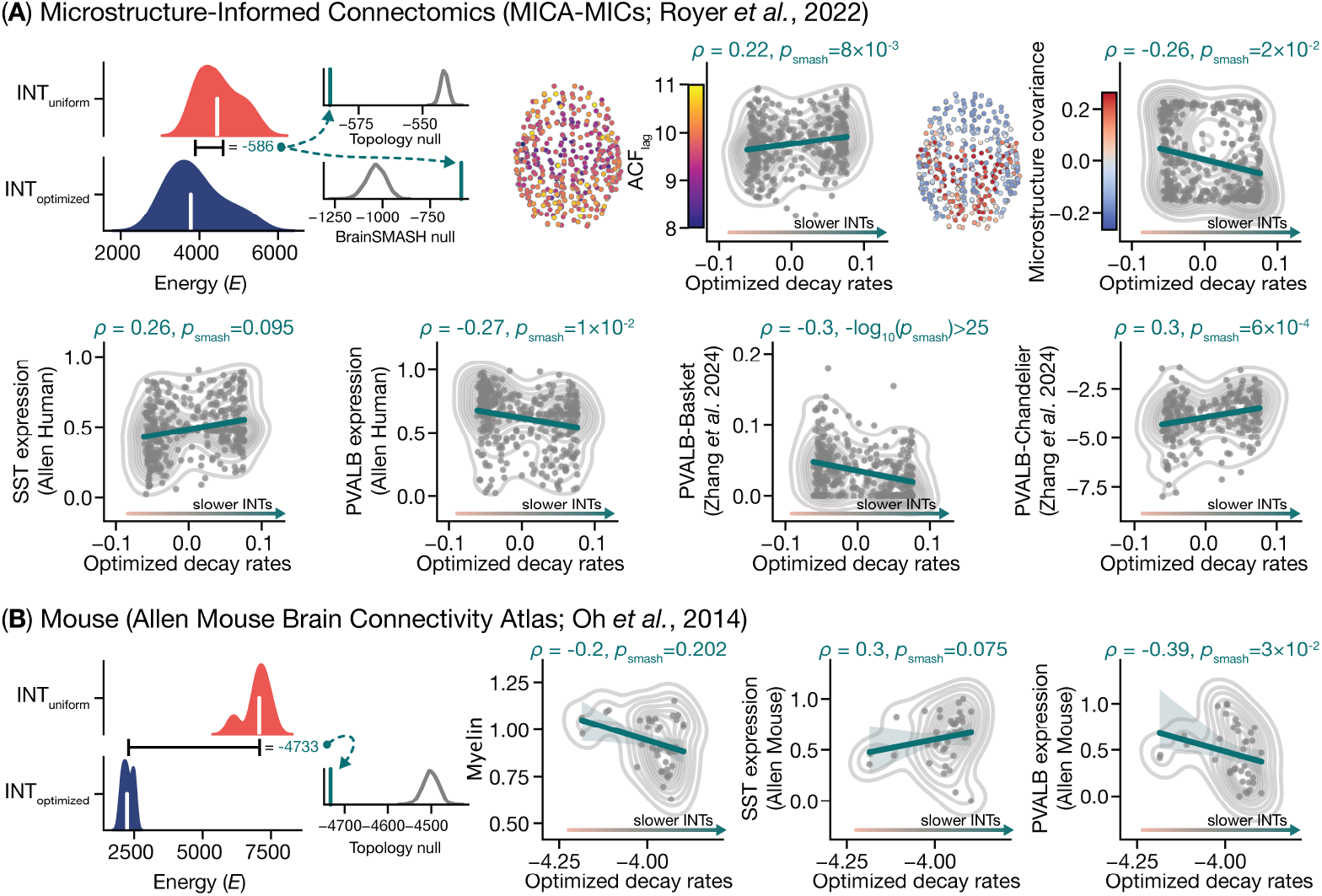
Results from the *INT*_*optimized*_ model replicate across datasets and generalize across species. We fit our model to two separate replication datasets. (**A**) Replication in the Microstructure-Informed Connectomics (MICA-MICs) human dataset [50]. We observed that our results replicated in an independent human dataset. (**B**) Replication in the Allen Mouse Brain Connectivity Atlas [47, 48]. We found that our results replicated in the mouse connectome, demonstrating that they generalized across species.

When examining the directed mouse connectome, we found that control energy was lower for the *INT*_*optimized*_ model compared to the *INT*_*uniform*_ model, and that this effect was larger than expected under our topology null (Fig. 6B, left). Note, we could not run the brainSMASH null here because we used binary brain states to examine state transitions on the mouse connectome (see Materials and Methods). Second, using maps of myelin as well as SST and PVALB genes extracted from the mouse isocortex [5], we observed the same set of correlations with optimized decay rates as those found in the human connectome (Fig. 6B, right). Note, while two of these correlations were not significant under our brainSMASH null, their signs and effect sizes were highly consistent with those observed in the human connectomes, suggesting that the lack of significance may have been a power issue. These results are consistent with prior work demonstrating correspondence between expression of inhibitory interneuron genes and multi-modal brain structure in the mouse brain [5, 75].

The results presented in this section demonstrate that the key outputs from the *INT*_*optimized*_ model replicate across multiple datasets and are conserved across species. Critically, our findings demonstrate convergent multi-scale linkages between brain structure and function across humans and mice. In both organisms, genes that track diverging inhibitory processes across the cortex—and that are thought to underpin regions’ myelin content and inter-connectivity—correlate consistently with regions’ INTs estimated from NCT. This result suggests that the exogenous control of large-scale brain dynamics from the connectome is rooted in microscale cellular processes, and that these effects are phylogenetically conserved.

### 2.7 Optimizing regions’ internal dynamics leads to improved prediction of cognition

Above, we demonstrated that our *INT*_optimized_ model gives rise to more efficient state transitions (Fig. 2. Fig. 3) and yields interpretable maps of brain regions’ intrinsic dynamics that correlate with empirical INTs (Fig. 4), gene expression and cell-type distributions (Fig. 5). However, all of these analyses were performed on a group-averaged connectome. One of the key advantages of NCT is that it is highly amenable to fitting to single-subject data [23]. Thus, in our final analysis, we fit our NCT model to individual connectomes (see Materials and Methods). Briefly, we fit both our *INT*_uniform_ and *INT*_optimized_ models to 960 subject-specific HCP-YA connectomes and examined the correlation between subject-specific control energy and subjects’ scores on a battery of behavioral tasks (see Materials and Methods).

Using a cross-validated penalized regression model, we examined whether control energy from our *INT*_optimized_ model outperformed the *INT*_uniform_ model at out-of-sample prediction of behavior (see Materials and Methods). Across 100 repeats of 5-fold cross-validation, we found that the *INT*_optimized_ model outperformed the *INT*_uniform_ model across all behavioral measures, including their average (Fig. 7). Additionally, empirical null models (see Materials and Methods) revealed that prediction performance for both variants of control energy exceeded chance levels (Fig. 7, insets; all *ps <* 0.05). These results show that optimizing the internal decay parameters of our NCT model at the single-subject level leads to a significant improvement in the out-of-sample prediction of cognition in the HCP-YA.

**Figure 7.**
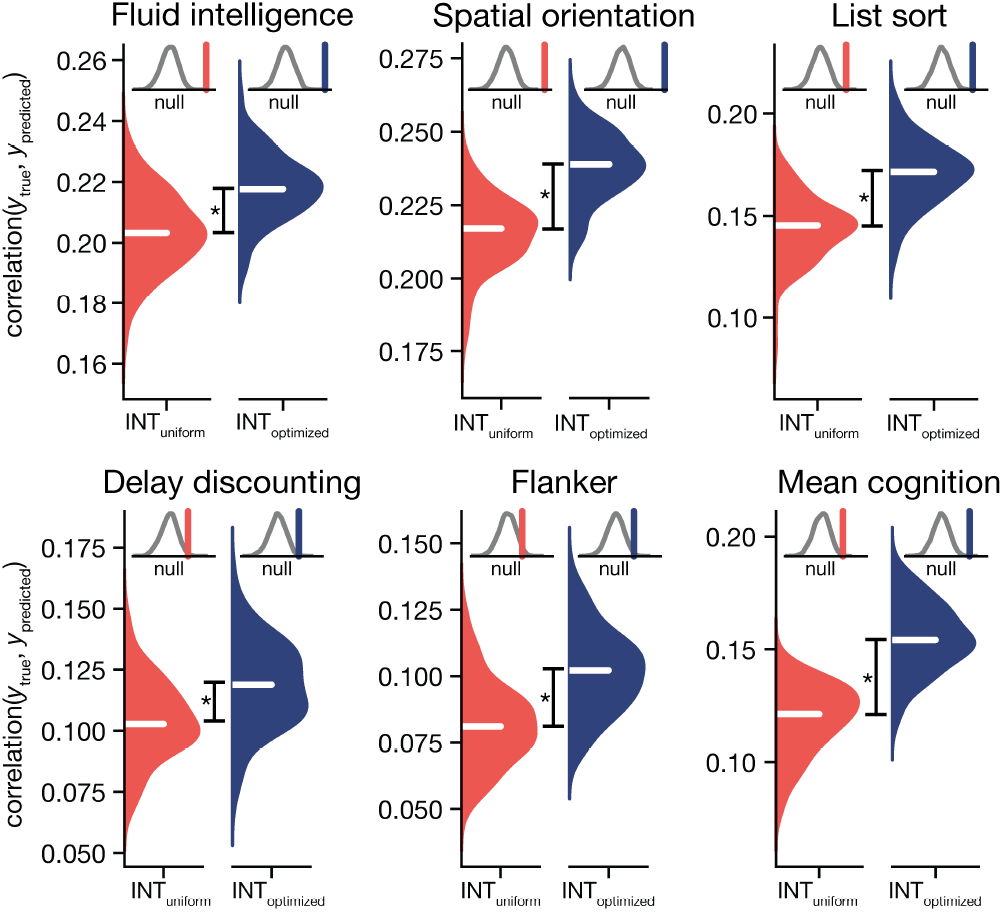
Subject-specific optimized control energy yields improved out-of-sample prediction of cognition. Results from a brain-based predictive modeling analysis that used subject-specific control energy to predict individuals’ cognition. We extracted subject-specific control energy from both *INT*_optimized_ and *INT*_uniform_ models and used those features as predictors in separate cross-validated penalized regression models (see Materials and Methods). Control energy from the *INT*_optimized_ model outperformed that of the *INT*_uniform_ model at predicting subjects’ cognitive scores. Permutation testing revealed that energy from both models performed above chance levels (insets). *= *p <* 0.05.

## 3 Discussion

We developed an extension to NCT that enabled us to model brain regions’ intrinsic neural timescales (INTs) from the structural connectome. Our approach corrects a long-held assumption embedded within the NCT framework: that brain regions’ intrinsic dynamics are fixed and homogeneous across the brain. This assumption has limited the biophysical realism of NCT, in turn hampering its application to neuroscience. Here, we correct this biologically implausible constraint by providing a data-driven approach to uncovering whole-brain maps of INTs that improve structure-function coupling. We found that model-based INTs reduced the control energy associated with transitioning between fMRI activity states and enabled the efficient completion of these transitions from fewer control nodes. We also found that model-based INTs correlated robustly with empirically-measured INTs as well as diverse measures of gene expression and cell-type densities measured *ex vivo*. Next, fitting our model to subject-specific data, we found that model-based INTs improved the out-of-sample prediction of behavior in a sample of young adults. Finally, we found that our key results replicated across two human datasets as well as in the mouse brain, and were robust to choice of brain state resolution (see Supplementary Materials). This work provides an intuitive transition- and subject-specific way of using NCT to infer measures of microscale neurobiology *in vivo*.

### 3.1 Modeling regions’ intrinsic neural timescales reduces control energy

Here, we observed that modifying the self-inhibition parameters in NCT consistently reduces the control energy needed to move between brain activity states. In the standard implementation of NCT (*INT*_uniform_), control signals are tasked with controlling brain activity by jointly leveraging the diverse connectivity between nodes as well as the fixed internal dynamics within nodes [23, 51, 52]. By contrast, our approach enables control signals to leverage both diverse extrinsic connectivity and diverse intrinsic dynamics to control brain activity. Notably, unlike the connectivity between nodes, these diverse intrinsic dynamics are not set as inputs, rather they are tunable by NCT during model fitting. In turn, this added parameter flexibility allows the *INT*_optimized_ model to find more efficient state transitions by uncovering, and then harnessing, transition-specific regional dynamics. We found that this added model flexibility yielded reduced control energy for the *INT*_optimized_ model, indicating that our approach narrowed the gap between brain structure and function using NCT. This observation is consistent with previous studies documenting increased structure-function coupling in network models that embrace more complexity in their assumptions about neural signaling [10, 13, 76].

We found that the difference in control energy between the two models was driven jointly by the topology of the connectome and the spatial patterning of the state transitions. Specifically, using a pair of null models [56, 57], we observed that the effects of connectome topology and the state transitions were in tension with one another. While control energy was always lower for the *INT*_optimized_ model compared to the *INT*_uniform_ model, the magnitude of this reduction was significantly larger than expected when compared to a connectome with edges that were rewired, and significantly smaller than expected compared to when the target states were permuted. This *tug of war* effect demonstrates that uncovering INTs using our approach required balancing the competing influences of connectome topology and the spatial properties of the brain states. This effect delivers two key insights. First, from a methodological perspective, this effect shows that the outputs of our algorithm were not a trivial consequence of either of the key model inputs (i.e., the connectome or the brain states). Second, this effect suggests that regions’ INTs exhibit both a stable and a flexible component. The stable component varies systematically across the cortex [2], as well as throughout development [22], and likely explains similarities in INTs between rest and task contexts [77, 78]. Our results indicate that this stable component is tied to the topology of the human connectome. However, while regions’ INTs are tightly constrained by a relatively static connectome topology, they are also inherently dynamic, which allows them to flexibly respond to different contexts (i.e., state transitions). This result is consistent with previous work showing that while INTs exhibit similar global structure across rest and task conditions, they also show clear modulation in response to specific task demands [78, 79]. This flexible component has implications for studies seeking to integrate spatially-patterned features of neurobiology into dynamical systems models [67, 80, 81], and highlights that incorporating variable context-specific patterns of INTs may enable more accurate modeling of large-scale neural systems.

### 3.2 Controlling brain dynamics effectively from fewer brain regions

A current shortcoming in the application of NCT to neuroscience is that practitioners generally adopt a full control set [23]. This approach involves delivering control signals to all nodes of the connectome. We have used this approach extensively in our previous work [8, 29, 34, 43]. However, controlling all brain regions at once is inconsistent with how numerous neurostimulation techniques, including transcranial magnetic stimulation (TMS), transcranial direct current stimulation (tDCS), and deep brain stimulation (DBS), affect the brain. In each of these techniques, stimulation is delivered focally instead of globally as in NCT. The reason for modeling whole-brain stimulation in NCT is, in part, a practical one. As discussed in our recent protocol paper [23], controlling neural dynamics from very few nodes is often intractable in NCT, which in turn necessitates a relatively broad control set in order to guarantee a successful state transition. To address this gap, here—as well as in our previous work [43]—we developed a novel extension to NCT that allowed us to model state transitions successfully from relatively few control nodes. We found that the *INT*_optimized_ model enabled successful low-energy state transitions from fewer regions than were required for the *INT*_uniform_ model (Fig. 3). This result demonstrates that expanding the biophysical realism of NCT has the potential to close the gap between the historical application of NCT and one of its clinical use cases (modeling neurostimulation). Finally, we observed a similar effect in our recent work that used spatially-diffuse control signals applied to the *INT*_uniform_ model [43]. Given the independence of these NCT modifications, future work could consider combining spatially-diffuse control signals with the *INT*_optimized_ model to further enhance the clinical translatability of NCT.

### 3.3 Optimized INTs correlate with empirically-measured neural timescales, neurobiology, and cognition

The approach we present here finds node-level INTs for the connectome through data-driven optimization of NCT decay rates. This stands in contrast to prior work that extended the biophysical realism of NCT by modifying control parameters *a priori* using neurobiology [39, 41]. While both approaches are viable, a key advantage of our data-driven approach is that it enables us to uncover both transition-specific and subject-specific INTs, and to use them as proxies for measures of neurobiology that are otherwise unobservable *in vivo*.

Here, we found that optimized INTs varied across state transitions and that the dominant mode of this variation coupled tightly to empirically-measured INTs estimated from fMRI data. That is, weaker self-inhibition in our model, which corresponds to slower intrinsic timescales, correlated with longer resting-state INTs in the human connectome. This finding validates the interpretation of optimized self-inhibition parameters in our model as reflecting nodes’ INTs. Moreover, we showed that our optimized INTs correlated systematically with diverse markers of inhibitory interneurons, including expression of SST and PVALB genes from the AHBA [45] as well as the density of PVALB-expressing basket and chandelier cells [46]. Critically, our effects tracked known divergences in these interneuron markers consistently across the human and the mouse brain, illustrating that our model-based INTs track variation in genes and cell-types that separate INTs across the cortex [3, 5, 75]. Together, these results robustly validate our approach as a means of uncovering biologically-grounded whole-brain INTs directly from the connectome *in vivo*. In turn, this opens up the possibility to use NCT to estimate and study subject-specific variation in spatially-patterned context-specific INTs.

Leveraging this subject specificity, we fit our model separately to 960 subjects from the HCP-YA young adult sample. In doing so, we found that control energy from the *INT*_optimized_ model outperformed that of the *INT*_uniform_ model at predicting a diverse range of behavioral measures out-of-sample. In sum, these results indicate that our *INT*_optimized_ model is able to both uncover valid estimates of brain regions’ INTs and leverage them to improve the strength of brain-behavior relationships.

### 3.4 Future directions

This study gives rise to two key future directions. First, the effects of development on the human connectome, brain regions’ INTs, and NCT are all well documented [8, 22, 82–85]. For example, hallmark features of connectome topology, including the presence of hub nodes [83, 84] and modular structure [82], emerge systematically throughout development, shaping the relationship between brain structure and function [86]. Accordingly, control energy reduces systematically throughout childhood and into young adulthood [8, 31, 85]. Critically, these developmental effects occur in lockstep with spatially-patterned shifts to brain regions’ empirical INTs [22] as well as features of brain structure that are thought to proxy regions’ gene expression (e.g., intracortical myelin content [87]). Our approach has the potential to connect these entwined phenomena to better understand how the multi-scale relationship between brain structure and function emerges throughout development.

A second extension is to integrate the *INT*_optimized_ model with existing NCT extensions that also seek to improve the biophysical realism of the model. As discussed above, we recently implemented an extension to NCT that involved inputting spatially-diffuse control signals into each node [43]. Briefly, this approach challenges another long-held assumption of NCT that each control signal exclusively impacts a signal node; an assumption that is inconsistent with the spatial continuity of the cerebral cortex and its effects on dynamics [88]. Notably, the spatial control strategy implemented in our prior work [43] and the *INT*_optimized_ model developed here involve modifications to separate components of the NCT equations. Specifically, the spatial strategy involves modifying only the control inputs, *B*, while the *INT*_optimized_ model involves modifying only the system, *A*. As such, they provide complementary viewpoints on improving the biophysical realism of NCT, and in turn may be combined in future studies.

### 3.5 Limitations

This study has limitations. First, we validated our optimized INTs empirically using fMRI data, which has a relatively low temporal resolution that, in turn, limits its capacity to capture the full spectrum of regions’ intrinsic dynamics. Future studies should consider applying the *INT*_optimized_ model to high temporal resolution data, such as magnetoencephalography. Second, as with all NCT studies, the underlying dynamical model assumes that large-scale brain dynamics are relatively well captured by a simplified linear model (see [23] for extended discussion). While this assumption limits some of insights provided by NCT, it also enables NCT to fit straightforwardly to whole-brain connectomes and across large datasets; a benefit that makes NCT and other linear approaches well-suited to studying individual differences. Furthermore, in this study, the assumption of linear dynamics allows for relatively fast model training to uncover INTs. This is because solutions to NCT are closed-form.

### 3.6 Conclusions

Here, we developed a novel extension to NCT that allowed us to uncover whole-brain patterns of brain regions’ intrinsic neural timescales from the human connectome. These timescales varied across the brain as well as across state transitions, and were replicated in both human and mouse brains. They also correlated with empirically-measured neural timescales as well as diverse markers of inhibitory interneurons consistently across species. Finally, when applied to subject-specific connectomes, our approach improved brain-behavior correlations across a wide range of cognitive tasks. In sum, our approach extends the biophysical realism of NCT and increases its translational utility for the field of neuroscience.

## 4 Materials and Methods

### 4.1 Participants

In this study, we analyzed three separate datasets. Our primary dataset, the results for which are reported throughout the main text, was the Human Connectome Project Young Adult (HCP-YA) sample [49]. We also sought to replicate our primary results in a second human dataset. For this purpose, we used the Microstructure-Informed Connectomics (MICA-MICs) dataset [50]. Lastly, we examined whether our results generalized across species by examining data from the Allen Mouse Brain Connectivity Atlas [47, 48]. We describe each of these datasets, as well as our processing of them, below.

#### 4.1.1 Human Connectome Project

The HCP-YA dataset [49] consists of diffusion-weighted magnetic resonance imaging (dMRI) data, resting state functional MRI (rs-fMRI) data, and structural MRI (T1-weighted and T2-weighted) data from young adult subjects. The HCP-YA was approved by the Washington University Institutional Review Board and informed consent was obtained from all subjects. Here, we used these MRI data from 960 HCP-YA subjects (age = 28.7*±*3.72; 53% female).

##### Diffusion-weighted magnetic resonance imaging (dMRI)

Probabilistic tractography was run on each subject’s minimally preprocessed dMRI data using MRtrix [89]. We generated 10^7^ streamlines. The 400-region variant of the Schaefer parcellation [54] was used to extract structural connectivity (SC) weights. We normalized the raw streamline counts by dividing them by the sum of the region sizes of their respective node pairs. Region sizes were estimated using subject-specific versions of the parcellation file. Then, we log-normalized the weights. These size- and log-normalized SC weights were taken as adjacency matrices, *A*, and used for NCT analysis. Note that *A*_*ij*_ = 0 for *i* = *j*. In cases where a group-averaged *A* matrix was used for analysis, we averaged over the entries of individuals’ *A* matrices and thresholded using an edge consistency–based approach. Specifically, edges in the group-averaged *A* matrix were only retained if non-zero SC weights were present in at least 60% of participants’ *A* matrices [90]. If not, then weights were set to zero.

##### Functional magnetic resonance imaging (fMRI)

Minimally preprocessed fMRI data (rfMRI_REST1_LR) were parcellated using the 400-region variant of the Schaefer parcellation [54]. Then, additional nuisance regression was performed that comprised the global signal (GS), the backward temporal derivative of the GS, as well as the square terms of both. This procedure created regional rs-fMRI time series that were used to create brain activity states for NCT analysis (see Section 4.3).

##### T1-weighted and T2-weighted magnetic resonance imaging (T1w, T2w)

Following prior work [63], we used the ratio of T1w to T2w MRI data as an *in vivo* proxy of intracortical myelin content. Vertex-level myelin maps were downloaded from the HCP-YA database for each subject (/MNINonLinear/fsaverage_LR32k/[subj_id].MyelinMap_BC_MSMAll.32k_fs_LR.dscalar.nii). As above, these maps were parcellated using the 400-region variant of the Schaefer parcellation [54].

##### Behavior

We extracted behavioral scores from the following tasks: fluid intelligence (PMAT24_A_CR), spatial orientation (VSPLOT_TC), list sort (ListSort_Unadj), delay discounting (DDisc_AUC_40K), and the flanker task (Flanker_Unadj). Additionally, we averaged across these measures to generate a single composite score for cognition. We used these scores as dependent variables—alongside age and sex as nuisance covariates—in our brain-based predictive modeling analysis (see Section 4.9).

#### 4.1.2 Replication Dataset: MICA-MICs

We used the MICA-MICs dataset [50] to replicate our primary results in another human dataset. The MICA-MICs dataset consists of dMRI and rs-fMRI data, among other modalities, from 50 adult subjects (age 29.54*±*5.62 years; 23 women).

##### Diffusion-weighted Magnetic Resonance Image (dMRI)

dMRI data were processed using *MRtrix3* [89]. Briefly, dMRI were preprocessed using the FSL pipeline built into *MRtrix3* (dwifslpreproc), which includes eddy and TOPUP. Response functions for the spherical deconvolution were estimated using the dhollander algorithm. Fibre orientation distributions were estimated using multi-shell multi-tissue constrained spherical deconvolution. 10^7^ streamlines were generated using the iFOD2 and SIFT2 algorithms with dynamic seeding from the white matter. As above, structural connectivity was estimated using the 400-region variant of the Schaefer parcellation [54], edge weights were size- and log-normalized, and consistency-based thresholding was used to generate a group-averaged *A* matrix.

##### Functional Magnetic Resonance Image (fMRI)

rs-fMRI data were processed using *fMRIprep* [91, 92] and *XCP-D* [93]. Full details of these processing steps can be found in the *boilerplate* content included below in Section 4.9. As above, regional fMRI time series were extracted using the 400-region variant of the Schaefer parcellation [54].

#### 4.1.3 Replication Dataset: Allen Mouse Brain Connectivity Atlas

##### Directed mouse connectome

We studied a directed structural connectome obtained from the Allen Mouse Brain Connectivity Atlas [47, 48]. Whole-brain structural connectomes were constructed with 2 *×* 10^5^ voxels at a spatial resolution of 100 µm (see [47, 48] for more details). Voxels were assigned to regions (coarse structures) according to a 3-D Allen Mouse Brain Reference Atlas. Isocortex was further divided into 6 systems (auditory, lateral, medial, prefrontal, somatomotor, and visual) based on prior work that applied community detection to identify stable modules [94]. These 6 systems were used to define binary brain states for the purposes of NCT analysis (see [23] for a discussion on the difference between binary and non-binary brain states). Connection strengths were modeled for all source and target voxels using data from 428 anterograde tracing experiments in wild type C57BL/6J mice [48]. Normalized connection strengths were obtained by dividing the connection strengths by the source and target region sizes. Here, we retained only the 43 isocortical regions. This process resulted in a fully-connected directed structural connectome that we used as an *A* matrix for NCT analysis. Note, this is the same mouse connectome we used previously (see [23]).

##### Myelin

We extracted a regional map of the T1w/T2w ratio in the mouse isocortex from [5]. This map was matched to 40 of the 43 isocortical regions in the above connectome.

##### Gene expression

We extracted a regional maps of SST and PVALB gene expression in the mouse isocortex from [5]. As above, these maps were matched to 40 of the 43 isocortical regions in the above connectome.

### 4.2 Linear Optimal Control

We model the control dynamics of the brain as a linear time-invariant dynamical system according to

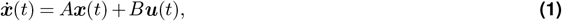

where ***x***(*t*) is an *N* -dimensional vector of activity across *N* brain regions, ***u***(*t*) is a *k*-dimensional vector of inputs, *A* is an *N × N* matrix of connectivity strength between brain regions, and *B* is an *N × k* matrix of connections from control inputs to brain regions. Here, we take *B* = *I* to be the identity matrix where *k* = *N* such that each region receives its own independent control input. This setup is known as a *uniform full control set* [23]. The goal of linear optimal control theory in NCT is to find the input ***u***(*t*) that transitions the system from an initial state ***x***_0_ to a final state ***x***_*T*_ while minimizing the following cost,

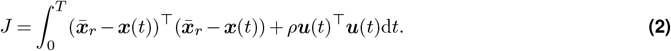

Here, 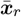 is the average reference state among a set of reference states, 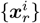.Throughout, we set 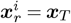 such that 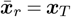,and call ***x***^*^(*t*) and ***u***^*^(*t*) the optimal trajectory and input, respectively. Note, 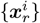 is user-modifiable and could be set to a variety of reference states (e.g.,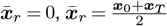). Note, we examined the effect of setting the reference state to a vector of zeros 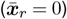,however this resulted in the model generating outputs optimized to simply *turn off* ***x***_0_, rather than completing the state transition. As such, we recommend setting 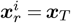.

To ensure that the system dynamics are stable, the original adjacency matrix derived from the connectome, *A*, is typically normalized such that the real component of its eigenvalues are negative [23]. This normalization is obtained by dividing *A* by a small constant plus its largest eigenvalue (thereby making all eigenvalues less than 1), and then subtracting the identity matrix,

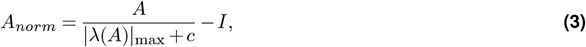

thereby ensuring all real components of the eigenvalues are less than 0. Here, this leads to regions’ internal dynamics being parameterized uniformly by a baseline decay rate of −1 (i.e., diag(*A*) = −1), an approach we refer to as *uniform intrinsic neural time scales* (*INT*_uniform_). While this uniform approach ensures stability, it fails to account for the heterogeneity in self-inhibition across brain regions [2, 19–22]. Here, we adjust regions’ decay rates to minimize the above cost function (Eq. 2).

We treat regions’ decay rates, ***d***, as learnable parameters given by a diagonal matrix *D* = diag(***d***) such that our new normalized adjacency matrix becomes

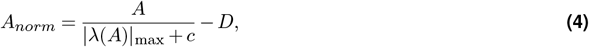

and we optimize *D* to minimize deviations between the state trajectory, ***x***(*t*), and the reference state, 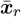,at the midpoint of the simulation, *t* = *T/*2. For this optimization, we implement optimal control in the automatic differentiation framework to perform backpropagation on ***d*** using the *PyTorch* library [95] first by normalizing the adjacency matrix (Eq. 4), then by numerically solving for the optimal trajectory ***x***^*^(*t*), and finally by penalizing deviations of the optimal trajectory at the midpoint. For the input matrix *B* = *I*, reference state 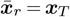,and the cost (Eq. 2) satisfying the boundary conditions of ***x***_0_ and ***x***_*T*_, the optimal trajectory can be found by taking derivatives of the control Hamiltonian

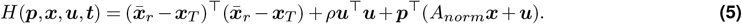

According to the Pontryagin minimum principle, if ***u***^*^(*t*) and ***x***^*^(*t*) are optimal inputs and trajectories, then there exists a time-varying Lagrange multiplier ***p***^*^(*t*) that satisfies 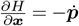,and 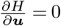,which can be written as

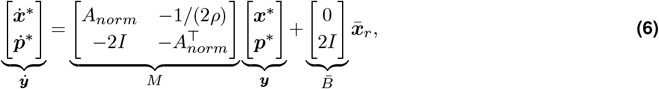

where we can write the full trajectory

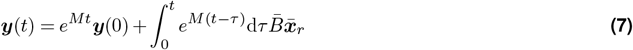

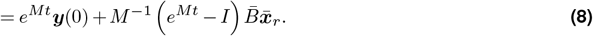

For 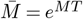 and 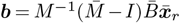 at the final time point, we use the initial and final states ***x***_0_, ***x***_*T*_ to solve for the initial condition of ***p*** by writing

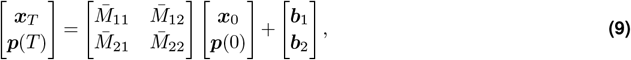

where 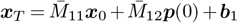,such that 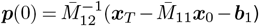.Using the initial conditions ***x***_0_, ***p***(0), we evolve the optimal state trajectory forward to the midpoint, compute the norm of the residual between the midpoint 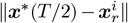,where 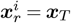 to yield the cost

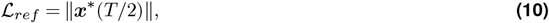

and backpropagate the error to the diagonal elements, ***d***.

Additionally, to achieve an optimized ***d*** that produces a normalized *A*_*norm*_ according to (Eq. 4) that is stable (i.e., all eigenvalues have negative real component), we add a second penalty to the eigenvalues *λ*_*i*_(*A*_*norm*_) to penalize their proximity to 0, such that

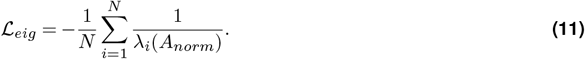

As the eigenvalues—which are initially all negative—approach 0, the loss will become large and diverge. Hence, as long as the learning rate is small enough, the eigenvalues should never vanish.

This optimization procedure comprises our *INT*_optimized_ model mentioned in the main text. We compared this *INT*_optimized_ model to a standard model where *A* was normalized according to Eq. 3. As noted above, we refer to this standard model as *INT*_uniform_.

### 4.3 Defining brain states

#### fMRI brain states

Following previous work [32, 39], we used temporal clustering to extract brain activity states from rs-fMRI data. We applied this approach to the HCP-YA and MICA-MICs datasets. First, for a given dataset, parcellated rs-fMRI time series were concatenated in time across all subjects, creating a data matrix, ***X***_*fmri*_, with *N* observations (rows) and *P* features (columns). Here, *N* is the number of time points in the rs-fMRI acquisition multiplied by the number of participants in the dataset and *P* is the number of brain regions in our parcellation (400). Next, we clustered ***X***_*fmri*_ in time using *k*-means clustering, sweeping through *k* = 2 − 19. To select the optimal *k* for our data, we examined the inertia of each clustering solution. At each *k*, inertia was estimated by calculating the distance between each data point (fMRI frame) and its respective centroid, squaring those distances, and summing across all distances separately for each centroid. Then, we compared the inertia of each clustering solution to the next and visually identified the point at which reductions to inertia plateaued. For HCP-YA, we chose *k* = 7 (see Fig. S1A). For MICA-MICs, we chose *k* = 5 (see Fig. S9E).

Note, we also reproduced our key findings for HCP-YA across *k* = 2 − 13 (see Fig. S6, Fig. S7, Fig. S8).

For *k* = 7 (*k* = 5 for MICA-MICs), we extracted the centroids of each cluster as brain states to be used for NCT analysis (i.e., ***x***_0_ and ***x***_*T*_). These brain states can be seen in Fig. 2A for HCP-YA. Each brain state represents a whole-brain map of regional co-activation that recurs over time throughout our rs-fMRI data. We characterized these brain states by (i) calculating their pairwise spatial correlations (*corr*(***x***_0_, ***x***_*T*_); see Fig. S1B) and (ii) quantifying their overlap with a canonical system-level organization of the brain (Fig. S1C) [53, 54]. Overlap was calculated by averaging centroid values within a set of binary brain masks that each represented one of seven canonical brain systems [53, 54].

#### Binary system-level brain states

We did not have access to neural activity data in the mouse brain. As such, we used the system-level organization discussed above in Section 4.1.3 to define binary brain states. This approach created 6 binary brain states comprising the auditory, lateral, medial, prefrontal, somatomotor, and visual systems.

### 4.4 Null models

In the main text, we discussed using a pair of null models to isolate and examine the effects of connectome topology as well as the brain states on our optimized decay rates. These null models are described in detail below. Using each null model, we re-fit our *INT*_optimized_ 5,000 times, generating a commensurate number of optimized decay rates for every state transition. This process produced a pair of null distributions for each state transition, one that examined the effect of connectome topology (Section 4.4.1) and another that examined the effect of the spatial-patterning of the brain states that comprised the transition (Section 4.4.2).

#### 4.4.1 Null network model

Our first null model consisted of randomly rewiring the connectome, each time preserving the spatial embedding of the network nodes as well as the network’s strength distribution [56]. Using this null network model, we produced 5,000 rewired networks derived from the group-averaged structural connectome used in the main text. Then, upon each rewired network, we re-fit our *INT*_optimized_ model for every state transition, producing 5,000 null optimized decay rates per transition.

#### 4.4.2 Brain state null model

Our second null model consists of randomly permuting the target state from each state transition. Note, here permuting the target also inherently also permutes the reference state, since these are one and the same. We performed this permutation using the Brain Surrogate Maps with Autocorrelated Spatial Heterogeneity (BrainSMASH) toolbox [57]. The spatial autocorrelation embedded in neuroimaging data can lead to inflated *p*-values in spatial correlation analyses and must be accounted for in the creation of null models. BrainSMASH addresses this by generating null brain maps that match the spatial autocorrelation properties of the input data. We used BrainSMASH to generate 5,000 spatial autocorrelation–preserving null maps of each brain state. Then, we re-fit our *INT*_optimized_ model for every state transition using each of these null brain maps as target/reference states. As above, this procedure produced 5,000 null optimized decay rates per transition.

### 4.5 Calculating control energy using random partial control sets

In the main text, we performed analyses that leveraged partial control sets (see Fig. 3). To conduct these analyses, random partial control sets were generated by first initializing the diagonals of *B* using zeros (diag(*B*) = 0) and then randomly setting a subset of those zeros to 1. The size of this random subset was determined by linearly sweeping through *k* = 5 to *k* = *N*_*system*_ *×* 0.5 to create 25 unique control set sizes. The estimation of control energy from a subset of nodes is a notoriously numerically poorly conditioned problem. As such, we estimated control energy using the most numerically stable algorithm known to us by computing the infinite-time controllability Gramian, *W*_*c*_, as a solution to the continuous-time Lyapunov equation. This approach does not require numerically challenging operations such as numerical integration and matrix exponentiation. We compute the control energy as the quadratic product of the target state with the inverse of the Gramian, and the estimated final state as 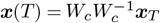.

### 4.6 Principal component analysis (PCA) of optimized decay rates

Our *INT*_optimized_ model produced a vector of regional optimized decay rates, ***d***, for every connectome and every state transition it was fit to. In the HCP-YA, we modeled state transitions between all possible pairwise combinations of seven brain states (see above). Thus, our analyses involved computing 42 state transitions, excluding self transitions, which in turn produced 42 sets of optimized decay rates. To characterize these decay rates, we compiled them into in a data matrix, ***X***_***d***_, wherein *N* state transitions were stored on the rows and *P* brain regions were stored on the columns. Then, we applied principal component analysis (PCA) to ***X***_***d***_, reducing the dimensionality of *N* to 5 PC maps that explained 91% of the variance in state transitions. This procedure gave us a compact set of brain maps that captured the dominant sources of variance in our optimized decay rates. Subsequently, we compared the first two PC maps with whole-brain maps of empirical intrinsic neural time scales (see Section 4.7), gene expression (see Section 4.8.1), and cell-type densities (see Section 4.8.2).

### 4.7 Quantifying empirical intrinsic neural time scales (INTs)

We used rs-fMRI data to quantify regions’ intrinsic neural time scales (INTs) empirically. INTs vary systematically across the cortex [96] and this variability is thought underpin the brain’s ability to process different types of information over different temporal receptive windows [2]. Following previous work [78], we estimated INTs by fitting an autocorrelation function (ACF) to each region’s rs-fMRI time series data and counting the number of time points it took for the ACF to decay to 0. We refer to this number as *ACF*_lag_ in the main text. Higher *ACF*_lag_ values indicate that it took more time points for the ACF to decay to 0, which we in turn interpret as slower fluctuations in the rs-fMRI data and thus slower INTs. By contrast, lower *ACF*_lag_ values indicate it took fewer time points for the ACF to decay to 0, which we interpret as faster fluctuations in the rs-fMRI data and thus faster INTs.

### 4.8 Whole-brain maps of neurobiology

As discussed in the main text, we examined the spatial coupling between our optimized decay rates and various maps of neurobiology that are thought underpin neural dynamics. Here, we outline the methods and tools we used to extract these maps.

#### 4.8.1 Gene expression from Allen Human Brain Atlas

We analyzed microarray data from the Allen Human Brain Atlas (AHBA), the full details of which can be found elsewhere [45]. Briefly, the AHBA includes normalized genome-wide microarray data (58,692 probes) for 3702 tissue samples taken from the cortex and subcortex of six post-mortem adult brains. We have used these data previously in combination with *in vivo* neuroimaging to bridge across multiple scales of neuroscience [97]. Here, using the abagen toolbox [59], we extracted parcellated whole-brain maps of gene expression following an established processing pipeline outlined in [60]. This process yielded a data matrix, ***X***_*gene*_, wherein expression of *N* genes were stored on the rows and *P* brain regions were stored on the columns. From ***X***_*gene*_, we extracted the rows corresponding to somatostatin (*SST*) and parvalbumin (*PVALB*) genes. Then, we correlated these rows with our optimized decay rates to examine the coupling being our model-based INTs and gene maps. We assigned p-values to these correlations using brainSMASH. Here, brainSMASH was used to generate 5,000 spatial autocorrelation–preserving maps of the first two principal components of optimized decay rates.

#### 4.8.2 Cell-type density maps from Zhang et al. [46]

While the AHBA affords unprecedented access to the spatially-patterned gene expression of large-scale brain systems, it is limited in its capacity to parse out specific cell types that express similar genes. One example of this is PVALB-expressing cells, which come in both Chandelier and Basket cell variants [74]. Addressing this gap, we recently developed an approach for imputing whole-brain cell-type density maps [46]. Briefly, we derived the gene expression signature for Chandelier and Basket PVALB-expressing cells—as well as twenty-two other cell types—from single-nucleus RNA-sequencing data [62] and then decomposed the fractions of each cell type from the total gene expression estimated from the AHBA microarray data (see [46] for more details). Using this approach, we extracted whole-brain maps of both PVALB-expressing Chandelier cells and PVALB-expressing Basket cells. Then, we correlated these maps with our optimized decay rates to examine the coupling being our model-based INTs and density of distinct PVALB-expressing cell types. As above, we assigned p-values to these correlations using brainSMASH.

### 4.9 Brain-based predictive modeling

As mentioned in the main text, we sought to link subject-specific control energy with individual differences in behavior using brain-based predictive modeling (BPM). To achieve this goal, we fit our *INT*_uniform_ and *INT*_optimized_ models to each subject’s structural connectome separately. For the *INT*_optimized_ model, this process generated 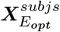, a 960 *×* 42 matrix of optimized control energy. Similarly, fitting the *INT*_uniform_ model created a 960 *×* 42 matrix, 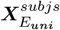,of standard control energy.

We iteratively fit penalized regression models using either 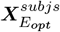 or 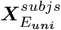 as predictors and a single measure of behavior as the output variable (see Section 4.1.1). Specifically, we used a cross-validated ridge regression model implemented in *scikit-learn* [98] with default parameters (*α* = 1) to predict participants’ behavioral scores (***y***). We assessed out-of-sample prediction performance using 5-fold cross-validation primarily scored by explained variance (*R*^2^) and secondarily scored by the correlation between the true ***y*** and predicted ***y***.

Regression models were trained using all columns of 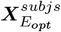 (or 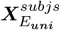)as input features and scoring metrics were each averaged across folds. We included age and sex as nuisance covariates. Nuisance covariates were controlled for by regressing their effect out of the predictors before predicting ***y***. Within each fold, nuisance covariates were fit to the training data and applied to the test data to prevent leakage. Finally, owing to evidence that prediction performance can be biased by the arbitrariness of a single split of the data [99], we repeated 5-fold cross-validation 100 times, each time with a different random 5-fold split. This process yielded a distribution of 100 mean *R*^2^ values and 100 mean correlations between true ***y*** and predicted ***y***.

Our above BPMs generated robust estimates of out-of-sample prediction performance that could be compared across predictors, but they did not examine whether prediction performance was itself significant. To test whether prediction performance exceeded chance levels, we compared point estimates of each of our scoring metrics—taken as the mean over the 100 values—to the distribution of values obtained from permuted data. Specifically, we subjected the point estimates of our scoring metrics to 5,000 random permutations, wherein the rows (i.e., participants) of ***y*** were randomly shuffled.

## Acknowledgements

JZK was supported by the Bethe/KIC/Wilkins, Eric and Wendy Schmidt AI in Science, and the Mong Neurotech postdoctoral fellowships. LP was supported by the National Institute Of Mental Health of the National Institutes of Health under Award Number R00MH127296. BL was supported by the National Institute Of Mental Health of the National Institutes of Health under Award Number R00MH127293.

## Code availability

Code can be found at: https://github.com/LindenParkesLab/nct_xr.

## Supplementary Materials

### Supplementary Figures

**Figure S1.**
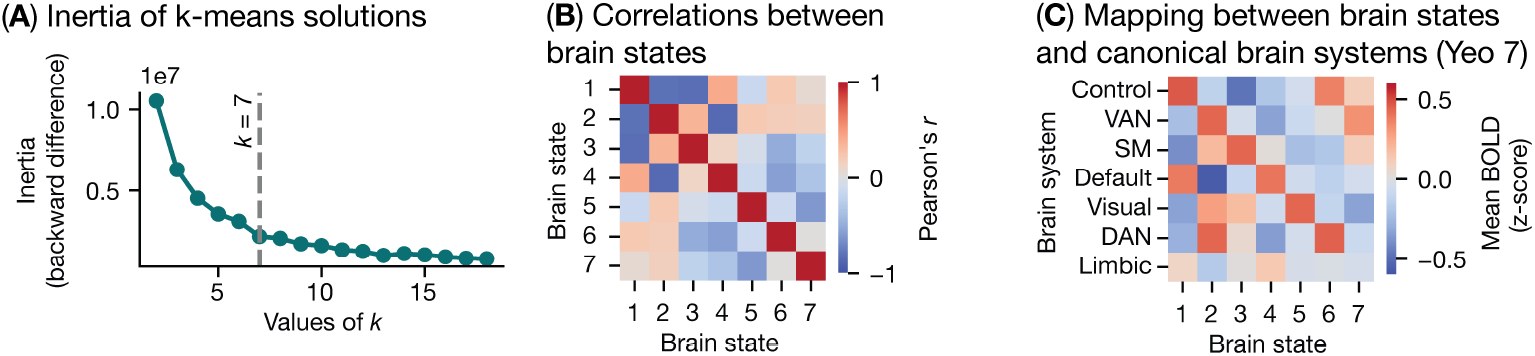
Characteristics of seven empirically-derived brain states using. (**A**) Inertia of *k*-means clustering solutions as a function of *k*. For each *k*, we calculated the inertia as the sum of squared distances between fMRI time points and their assigned cluster. Then, we examined the difference in inertia between *k* and *k* + 1 and determined that *k* = 7 was the optimal solution. (**B**) Pairwise spatial correlations between brain states. These correlations index brain states’ similarity. (**C**) Overlap between brain states and seven canonical brain systems [53]. Z-scored fMRI activity from each brain state was averaged over regions that comprised each canonical system. State 1 and 2 are mirror images of one another, with State 1 consisting of activation of the control and default networks as well as deactivation of the ventral attention network (VAN), somatomotor network (SM), visual system, and the dorsal attention network (DAN). By contrast, state 2 shows strong deactivation of the default network, and activation of the VAN, SM, visual system, and DAN. Thus, states 1 and 2 capture activity broadly cycling between the control/default networks and the rest of the brain, which is a common motif in resting-state brain dynamics. States 3 through 6 map predominantly onto the SM, default network, visual system, and the DAN, respectively. Finally, state 7 jointly captures the control, VAN, and SM. No state specifically captures activation of the limbic system.

**Figure S2.**
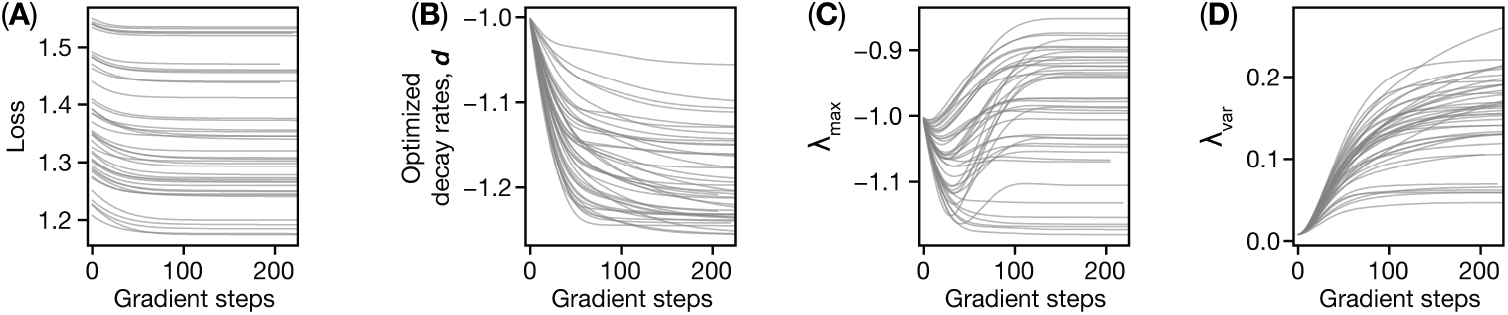
Parameters from the *INT*_optimized_ model plotted as a function of training. In each subplot, gray lines represent different state transitions. (**A**) Optimization loss as a function training. Here, loss is a combination of the reference state cost function (*ℒ* _*ref*_) and the eigenvalue cost function (*ℒ* _*eig*_); see Materials and Methods for more details. (**B**) Optimized decay rates, ***d***, plotted as function of training averaged over brain regions. (**C**) The maximum eigenvalue of the optimized connectome plotted as a function of training. (**D**) Variance in the eigenvalues of the optimized connectome plotted as a function of training. In **C** and **D**, eigenvalues were calculated by setting the diagonal of *A*_*norm*_ to ***d*** at each gradient step. The results presented here reveal the following insights. First, that our *INT*_optimized_ model stabilized quickly and early during training, with loss for each state transition reaching a plateau after ∼100 gradient steps in **A**. Second, that our model increased the overall strength of nodes’ self-inhibition throughout training (i.e., more negative optimized decay rates in **B**). Third, that the dominant mode of system dynamics remained inclined towards self-inhibition (i.e., negative *λ*_max_ in **C**) despite modifying nodes’ intrinsic dynamics. Finally, that the variability in system dynamics increased throughout training (i.e., greater *λ*_var_ in **D**). This last result demonstrates that our *INT*_optimized_ model gave rise to more diverse system-wide neural dynamics.

**Figure S3.**
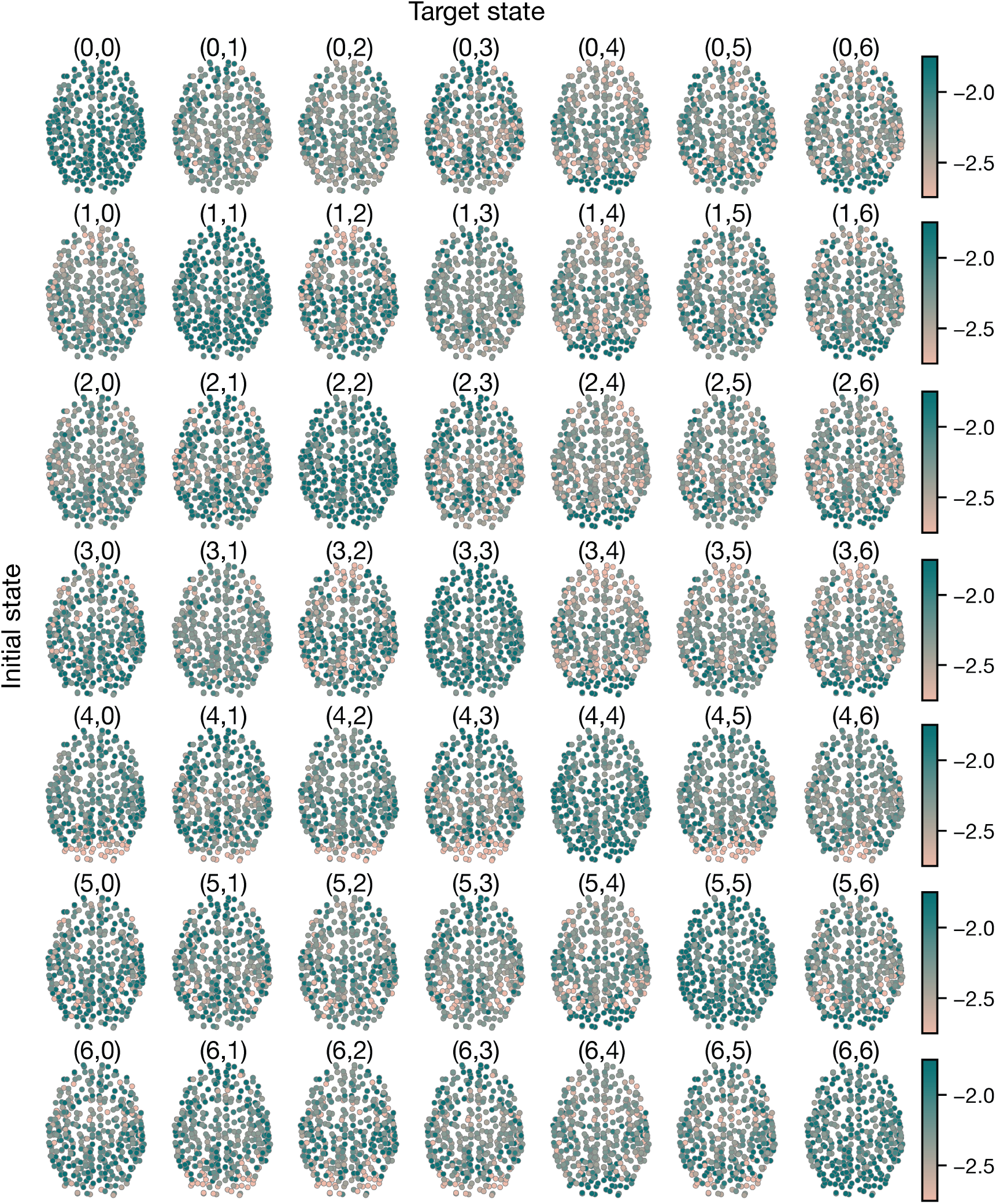
Optimized decay rates from the *INT*_optimized_ model for each state transition.

**Figure S4.**
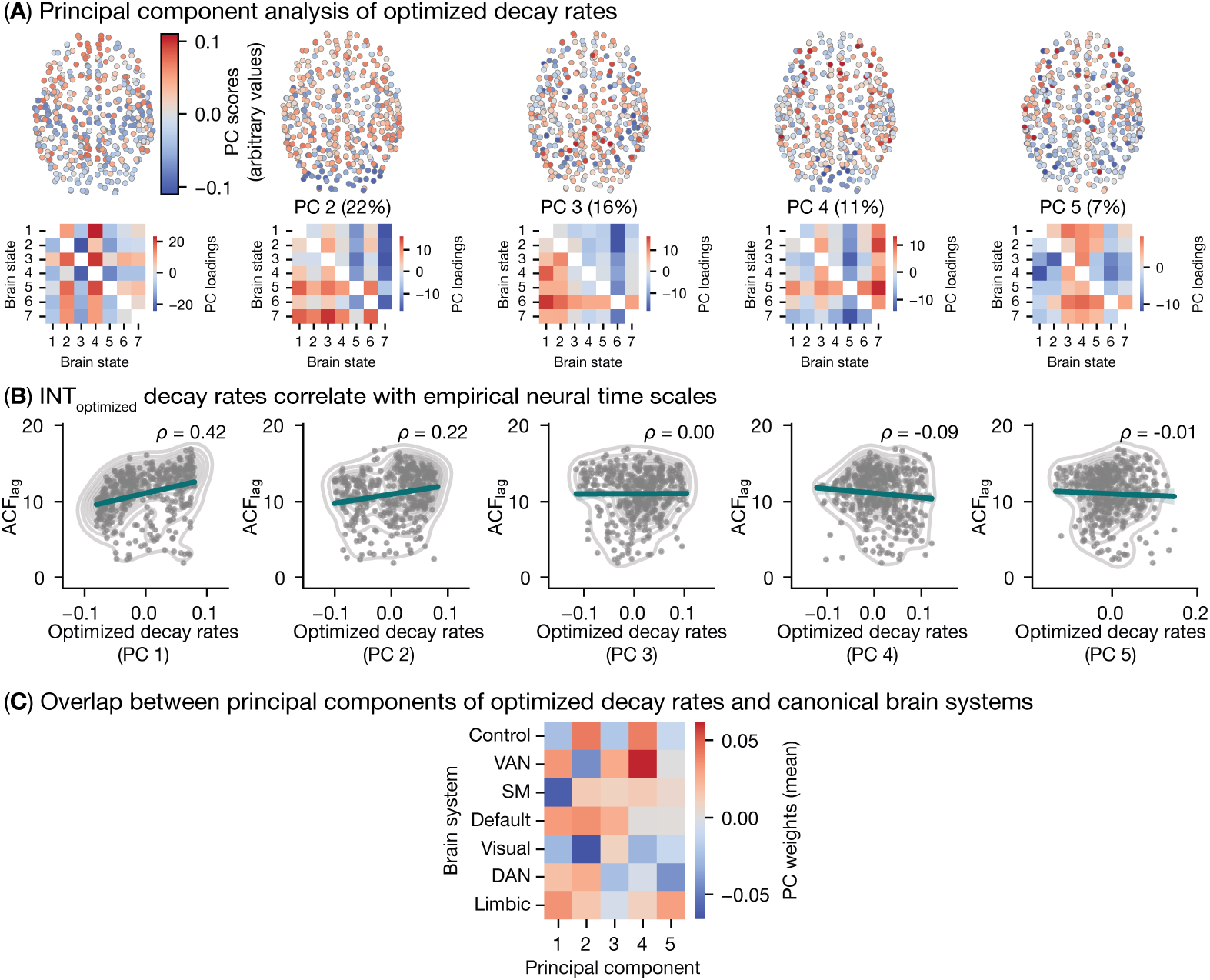
Model-based intrinsic neural time scales (INTs) correlate with empirically-measured INTs. (**A**) We used principal component analysis (PCA) to characterize variance in optimized decay rates over all state transitions. We extracted 5 PCs that explained 91% of the variance in optimized decay rates. (**B**) Correlations between PCs of optimized decay rates and empirically-measured INTs, as indexed by *ACF*_lag_ (see main text). (**C**) Overlap between PCs of optimized decay rates and canonical brain systems. Here, we plot the PC scores for each PC (*x* -axis) averaged over regions that comprise seven canonical brain systems (*y* -axis).

**Figure S5.**
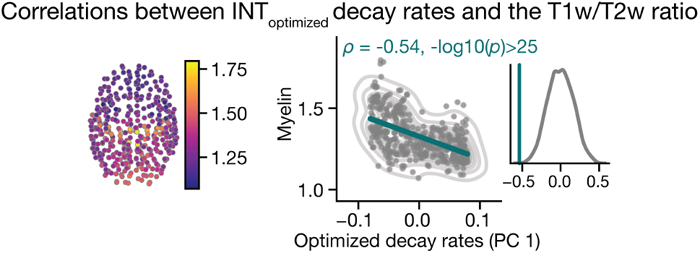
Model-based intrinsic neural time scales correlate with intracortical myelin content. Correlations between PC 1 of optimized decay rates (taken from Fig. 4) and the T1w/T2w ratio. The p-value was assigned using brainSMASH.

**Figure S6.**
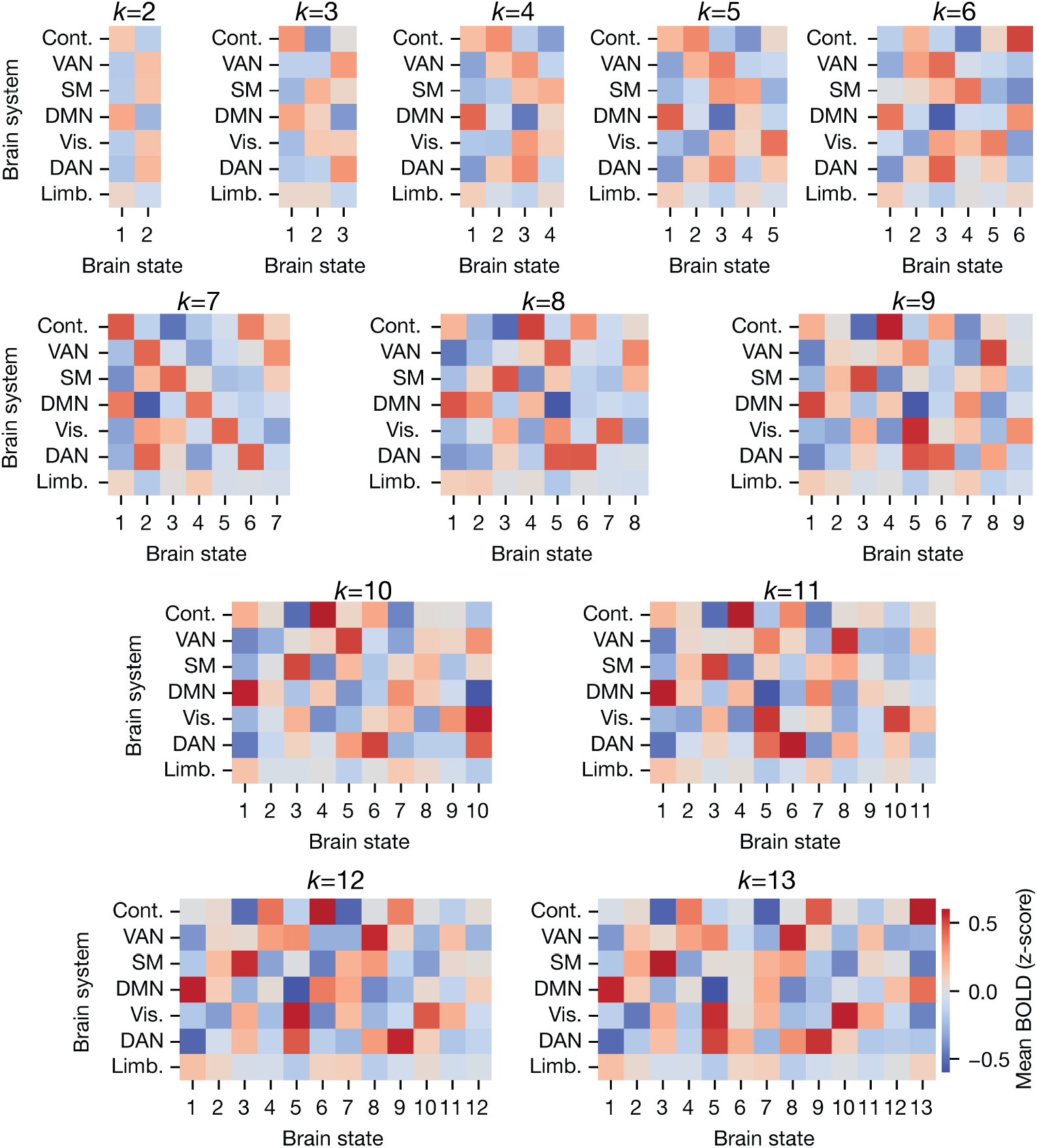
Overlap between seven canonical brain systems (rows) and brain states (columns) as a function of *k*.

**Figure S7.**
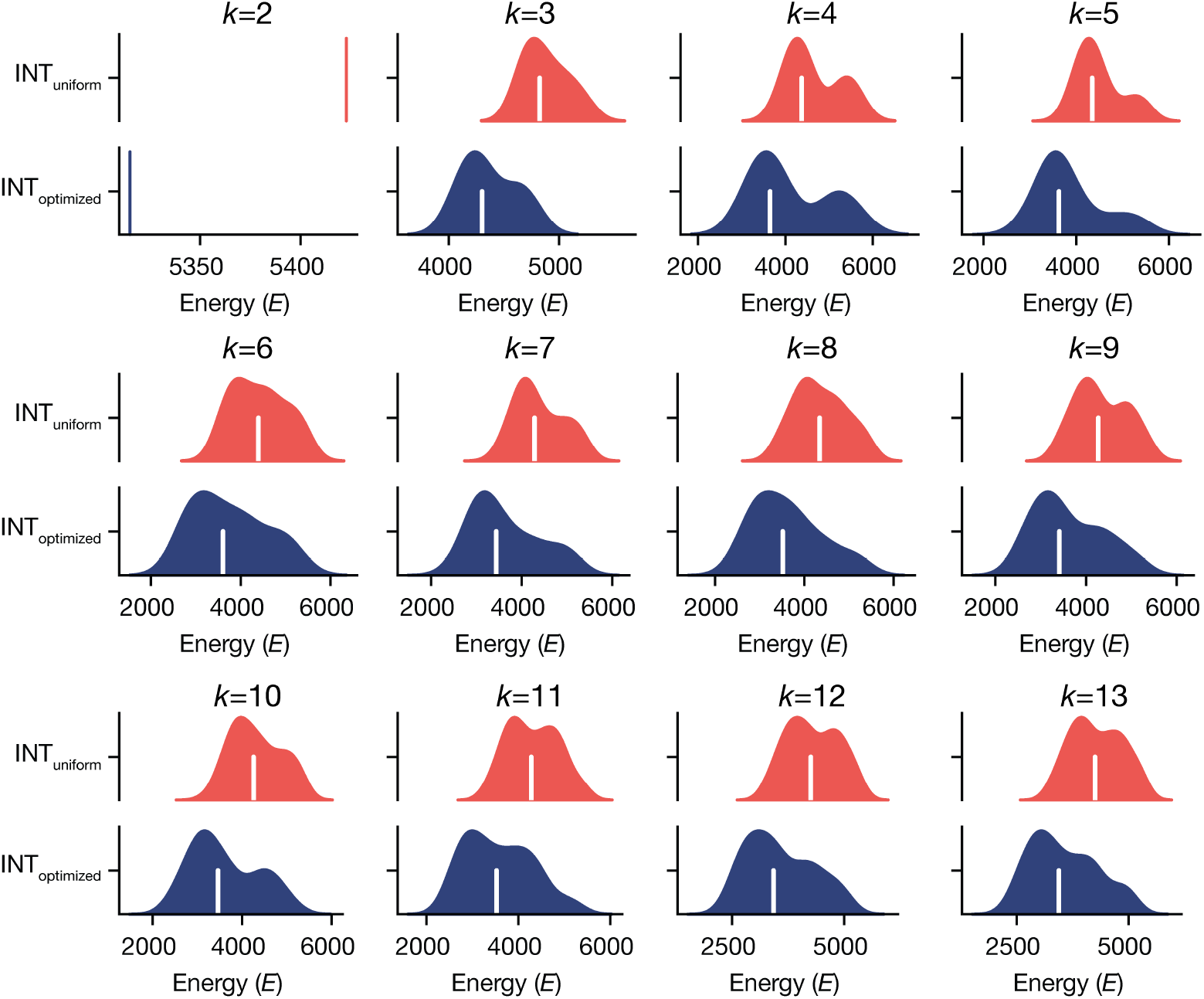
Control energy for the *INT*_uniform_ and *INT*_optimized_ models plotted as a function of *k*.

**Figure S8.**
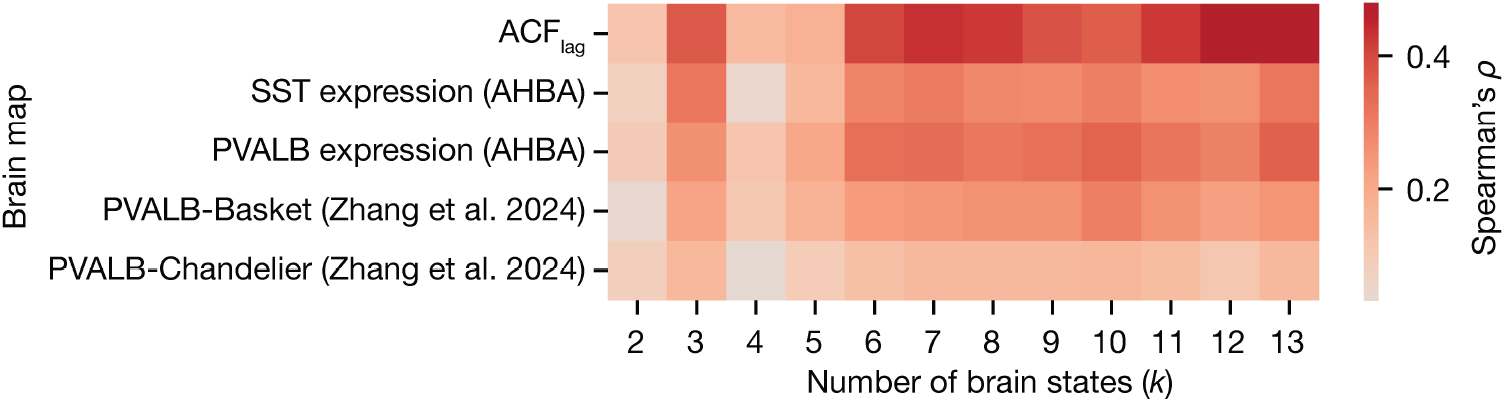
Correlations between optimized decay rates and empirical brain maps as a function of *k*. For each clustering solution, *k*, we extracted the first principal component of the optimized decay rates across all state transitions. Then, we correlated that principal component map with the five empirical brains maps discussed in the main text. Note, owing to the fact that the axes of a given principal component analysis are arbitrary, here we plot the absolute Spearman correlation coefficients.

**Figure S9.**
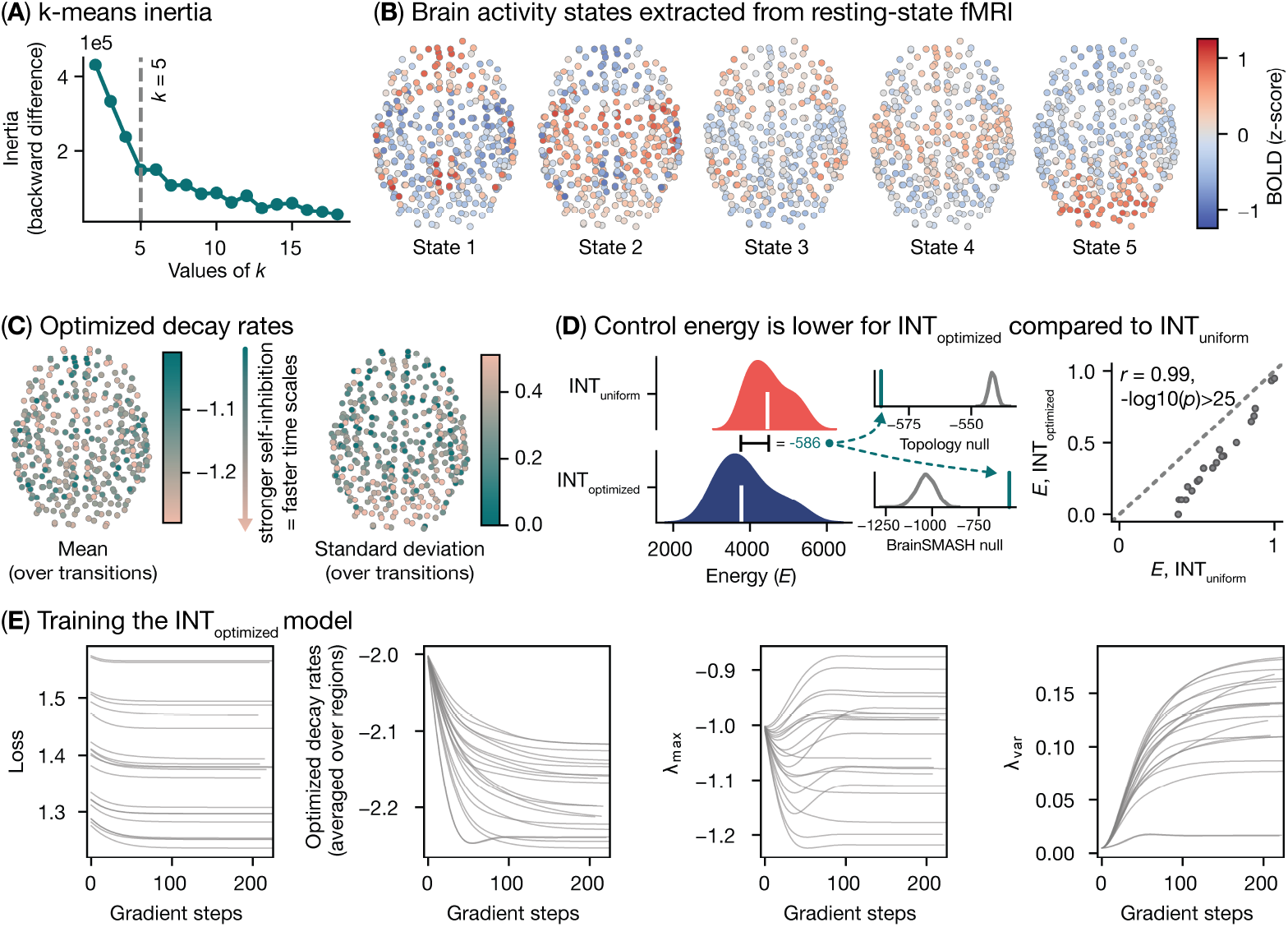
Replication dataset: MICA-MICs.

### Sensitivity analysis

Below, we reproduce several of our key findings and figures in a series of replication analyses outlined below.

### Different clustering solutions

In the main text, we presented our primary results by modeling state transitions between seven empirical brain states. These states were extracted by applying *k*-means clustering to rs-fMRI time series data with *k* = 7. Here, we replicate our analyses across *k* = 2 − 13. Figure S6 reproduces Figure S1C as a function of *k* by illustrating how brain states from each clustering solution aligned with canonical brain systems [53]. Figure S7 reproduces Figure 2C as a function of *k*, illustrating that control energy was lower for *INT*_optimized_ compared to *INT*_uniform_ across *k* = 2 − 13.

Finally, Figure S8 reproduces Figure 4 and Figure 5 as a function of *k* by showing that the first PC of optimized decay rates correlates consistently with *ACF*_lag_, gene expression, and cell-type densities.

### Replication datasets

Below, we replicate our main findings using our replication dataset (see Section 4.1.2).

Using the MICA-MICs dataset [50], we reproduced Fig. 2 and Fig. S2 (Fig. S9). Specifically, we extracted brain states using *k*-means clustering applied to the MICA-MICs rs-fMRI data and fit the *INT*_uniform_ and *INT*_optimized_ models to the *k* = 5 solution. We found that our results replicated, with the *INT*_optimized_ model yielding lower control energy across all state transitions. These replication efforts demonstrate that our findings were robust to differences in sample characteristics, data acquisition, data processing, and state definition.

### Boilerplate material

#### 4.9.1 fMRIprep

Some of the results included in this manuscript come from preprocessing performed using *fMRIPrep* 24.1.0 (Esteban et al. [100]; Esteban et al. [101]; RRID:SCR_016216), which is based on *Nipype* 1.8.6 (Gorgolewski et al. [102]; Gorgolewski et al. [103]; RRID:SCR_002502).

### Preprocessing of B0 inhomogeneity mappings

A total of 1 fieldmaps were found available within the input BIDS structure for this particular subject. A *B0*-nonuniformity map (or *fieldmap*) was estimated based on two (or more) echo-planar imaging (EPI) references with topup (Andersson et al. [104]; FSL None).

### Anatomical data preprocessing

A total of 2 T1-weighted (T1w) images were found within the input BIDS dataset. Each T1w image was corrected for intensity non-uniformity (INU) with `N4BiasFieldCorrection` [105], distributed with ANTs 2.5.3 [106, RRID:SCR_004757]. The T1w-reference was then skull-stripped with a *Nipype* implementation of the `antsBrainExtraction.sh` workflow (from ANTs), using OASIS30ANTs as target template. Brain tissue segmentation of cerebrospinal fluid (CSF), white-matter (WM) and gray-matter (GM) was performed on the brain-extracted T1w using `fast` [FSL (version unknown), RRID:SCR_002823, 107]. An anatomical T1w-reference map was computed after registration of 2 <module ‘nipype.interfaces.image’ from ‘/opt/conda/envs/fmriprep/lib/python3.11/site-packages/nipype/interfaces/image.py’> images (after INU-correction) using `mri_robust_template` [FreeSurfer 7.3.2, 108]. Brain surfaces were reconstructed using `recon-all` [FreeSurfer 7.3.2, RRID:SCR_001847, 109], and the brain mask estimated previously was refined with a custom variation of the method to reconcile ANTs-derived and FreeSurfer-derived segmentations of the cortical gray-matter of Mindboggle [RRID:SCR_002438, 110]. Volume-based spatial normalization to two standard spaces (MNI152NLin6Asym, MNI152NLin2009cAsym) was performed through nonlinear registration with `antsRegistration` (ANTs 2.5.3), using brain-extracted versions of both T1w reference and the T1w template. The following templates were were selected for spatial normalization and accessed with *TemplateFlow* [24.2.0, 111]: *FSL’s MNI ICBM 152 non-linear 6th Generation Asymmetric Average Brain Stereotaxic Registration Model* [Evans et al. [112], RRID:SCR_002823; TemplateFlow ID: MNI152NLin6Asym], *ICBM 152 Nonlinear Asymmetrical template version 2009c* [Fonov et al. [113], RRID:SCR_008796; TemplateFlow ID: MNI152NLin2009cAsym].

### Functional data preprocessing

For each of the 1 BOLD runs found per subject (across all tasks and sessions), the following preprocessing was performed. First, a reference volume was generated, using a custom methodology of *fMRIPrep*, for use in head motion correction. Head-motion parameters with respect to the BOLD reference (transformation matrices, and six corresponding rotation and translation parameters) are estimated before any spatiotemporal filtering using `mcflirt` [FSL, 114]. The estimated *fieldmap* was then aligned with rigid-registration to the target EPI (echo-planar imaging) reference run. The field coefficients were mapped on to the reference EPI using the transform. The BOLD reference was then co-registered to the T1w reference using `bbregister` (FreeSurfer) which implements boundary-based registration [115]. Co-registration was configured with six degrees of freedom. Several confounding time-series were calculated based on the *preprocessed BOLD*: framewise displacement (FD), DVARS and three region-wise global signals. FD was computed using two formulations following Power (absolute sum of relative motions, Power et al. [116]) and Jenkinson (relative root mean square displacement between affines, Jenkinson et al. [114]). FD and DVARS are calculated for each functional run, both using their implementations in *Nipype* [following the definitions by 116]. The three global signals are extracted within the CSF, the WM, and the whole-brain masks. Additionally, a set of physiological regressors were extracted to allow for component-based noise correction [*CompCor*, 117]. Principal components are estimated after high-pass filtering the *preprocessed BOLD* time-series (using a discrete cosine filter with 128s cut-off) for the two *CompCor* variants: temporal (tCompCor) and anatomical (aCompCor). tCompCor components are then calculated from the top 2% variable voxels within the brain mask. For aCompCor, three probabilistic masks (CSF, WM and combined CSF+WM) are generated in anatomical space. The implementation differs from that of Behzadi et al. in that instead of eroding the masks by 2 pixels on BOLD space, a mask of pixels that likely contain a volume fraction of GM is subtracted from the aCompCor masks. This mask is obtained by dilating a GM mask extracted from the FreeSurfer’s *aseg* segmentation, and it ensures components are not extracted from voxels containing a minimal fraction of GM. Finally, these masks are resampled into BOLD space and binarized by thresholding at 0.99 (as in the original implementation). Components are also calculated separately within the WM and CSF masks. For each CompCor decomposition, the *k* components with the largest singular values are retained, such that the retained components’ time series are sufficient to explain 50 percent of variance across the nuisance mask (CSF, WM, combined, or temporal). The remaining components are dropped from consideration. The head-motion estimates calculated in the correction step were also placed within the corresponding confounds file. The confound time series derived from head motion estimates and global signals were expanded with the inclusion of temporal derivatives and quadratic terms for each [118]. Frames that exceeded a threshold of 0.5 mm FD or 1.5 standardized DVARS were annotated as motion outliers. Additional nuisance timeseries are calculated by means of principal components analysis of the signal found within a thin band (*crown*) of voxels around the edge of the brain, as proposed by [119]. All resamplings can be performed with *a single interpolation step* by composing all the pertinent transformations (i.e. head-motion transform matrices, susceptibility distortion correction when available, and co-registrations to anatomical and output spaces). Gridded (volumetric) resamplings were performed using `nitransforms`, configured with cubic B-spline interpolation.

Many internal operations of *fMRIPrep* use *Nilearn* 0.10.4 [120, RRID:SCR_001362], mostly within the functional processing workflow. For more details of the pipeline, see the section corresponding to workflows in *fMRIPrep*’s documentation.

### Post-processing of fmriprep outputs

The eXtensible Connectivity Pipeline-DCAN (XCP-D) [121–123] was used to post-process the outputs of *fMRIPrep* version 24.1.1 [124, 125, RRID:SCR_016216]. XCP-D was built with *Nipype* version 1.8.6 [102, RRID:SCR_002502].

### Segmentations

The following atlases were used in the workflow: the Schaefer Supplemented with Subcortical Structures (4S) atlas [126–130] at 3 different resolutions (156, 256, and 456 parcels), the Schaefer1007 atlas, the Schaefer2007 atlas, and the Schaefer4007 atlas. Each atlas was warped to the same space and resolution as the BOLD file.

### Anatomical data

Native-space T1w images were transformed to MNI152NLin2009cAsym space at 1 mm3 resolution.

### Functional data

For each of the one BOLD runs found per subject (across all tasks and sessions), the following post-processing was performed.

In total, 36 nuisance regressors were selected from the preprocessing confounds, according to the ‘36P’ strategy. These nuisance regressors included six motion parameters, mean global signal, mean white matter signal, mean cerebrospinal fluid signal with their temporal derivatives, and quadratic expansion of six motion parameters, tissue signals and their temporal derivatives [123, 131].

The BOLD data were despiked with *AFNI*’s *3dDespike*.

Nuisance regressors were regressed from the BOLD data using a denoising method based on *Nilearn*’s approach. The timeseries were band-pass filtered using a(n) second-order Butterworth filter, in order to retain signals between 0.01-0.08 Hz. The same filter was applied to the confounds. The resulting time series were then denoised using linear regression. The denoised BOLD was smoothed using *Nilearn* with a Gaussian kernel (FWHM=6 mm).

The amplitude of low-frequency fluctuation (ALFF) [132] was computed by transforming the mean-centered, standard deviation-normalized, denoised BOLD time series to the frequency domain. The power spectrum was computed within the 0.01-0.08 Hz frequency band and the mean square root of the power spectrum was calculated at each voxel to yield voxel-wise ALFF measures. The resulting ALFF values were then multiplied by the standard deviation of the denoised BOLD time series to retain the original scaling. The ALFF maps were smoothed with Nilearn using a Gaussian kernel (FWHM=6 mm). Regional homogeneity (ReHo) [133] was computed with neighborhood voxels using *AFNI*’s *3dReHo* [134].

Processed functional timeseries were extracted from the residual BOLD signal with *Nilearn’s NiftiLabelsMasker* for the atlases. Corresponding pair-wise functional connectivity between all regions was computed for each atlas, which was operationalized as the Pearson’s correlation of each parcel’s unsmoothed timeseries. In cases of partial coverage, uncovered voxels (values of all zeros or NaNs) were either ignored (when the parcel had >50.0% coverage) or were set to zero (when the parcel had <50.0% coverage).

Many internal operations of *XCP-D* use *AFNI* [135, 136], *ANTS* [137], *TemplateFlow* version 24.2.2 [138], *matplotlib* version 3.9.2 [139], *Nibabel* version 5.3.0 [140], *Nilearn* version 0.10.4 [141], *numpy* version 2.1.2 [142], *pybids* version 0.17.2 [143], and *scipy* version 1.14.1 [144]. For more details, see the *XCP-D* website (https://xcp-d.readthedocs.io).

### Copyright Waiver

The above methods description text was automatically generated by *XCP-D* with the express intention that users should copy and paste this text into their manuscripts *unchanged*. It is released under the CC0 license.

